# Dynamics of diffusive cell signaling relays

**DOI:** 10.1101/2019.12.27.887273

**Authors:** Paul B. Dieterle, Jiseon Min, Daniel Irimia, Ariel Amir

**Affiliations:** Department of Physics, Harvard University, Cambridge, MA 02138, USA; Department of Molecular and Cellular Biology, Harvard University, Cambridge, MA 02138, USA; BioMEMS Resource Center and Center for Surgery, Innovation and Bioengineering, Department of Surgery, Massachusetts General Hospital, Boston, MA 02114, USA; John A. Paulson School of Engineering and Applied Sciences, Harvard University, Cambridge, MA 02138, USA

## Abstract

Cells can communicate with each other by emitting diffusible signaling molecules into the surrounding environment. However, simple diffusion is slow. Even small molecules take hours to diffuse millimeters away from their source, making it difficult for thousands of cells to coordinate their activity over millimeters, as happens routinely during development and immune response. Moreover, simple diffusion creates shallow, Gaussian-tailed concentration profiles. Attempting to move up or down such shallow gradients – to chemotax – is a difficult task for cells, as they see only small spatial and temporal concentration changes. Here, we demonstrate that cells utilizing diffusive relays, in which the presence of one type of diffusible signaling molecule triggers participation in the emission of the same type of molecule, can propagate fast-traveling diffusive waves that give rise to steep chemical gradients. Our methods are general and capture the effects of dimensionality, cell density, signaling molecule degradation, pulsed emission, and cellular chemotaxis on the diffusive wave dynamics. We show that system dimensionality – the size and shape of the extracellular medium and the distribution of the cells within it – can have a particularly dramatic effect on wave initiation and asymptotic propagation, and that these dynamics are insensitive to the details of cellular activation. As an example, we show that neutrophil swarming experiments exhibit dynamical signatures consistent with the proposed signaling motif. Interpreted in the context of these experiments, our results provide insight into the utility of signaling relays in immune response.

---

Prototypical diffusive signaling – in which individual cells communicate with neighbors by releasing diffusible molecules into the extracellular medium – is a relatively slow process. Signaling molecules undergoing random walks in the extracellular medium have a root mean square displacement that grows like the square root of both the time since emission, *t*, and the signaling molecule diffusivity, *D*. It follows that the distance an individual cell can signal also grows like the square root of time. Thus, for thousands of cells coordinating actions over millimeters, simple diffusive signaling with small molecules (*D* ≈ 10^−10^ m^2^/s) takes hours. These length and times scales are incommensurate with observed behavior in developmental biology [1–3], immune response [4], and microbial consortia [5], in which cells exchanging diffusible molecules coordinate activity over millimeters in tens of minutes.

Indeed, when many cells collectively integrate environmental cues and participate in the signaling, they can propagate diffusive waves with a fixed speed, *v*, in the asymptotic limit. This effect and its analogs have long been studied in the context of excitable media [6–8] and observed in biological phenomena as diverse as natural cell signaling circuits [9–12], synthetic cell signaling circuits [5], apoptosis [2], range expansions [13–18], and development [1, 3, 19]. In this way, small groups of cells can transmit signals more quickly than simple diffusion allows by recruiting the help of their neighbors.

Less well-understood is how the propagation and initiation of such waves are affected by the dimensionalities of the cellular distribution and the diffusive environment – or even how to identify the system dimensionality – as previous work has largely assumed quasi-1D dynamics [11, 12, 20]. Also unclear are how robust the resulting signaling dynamics are to underlying biological details.

Here, through a comprehensive study of single-component relays — in which cells measure the local concentration of a signaling molecule and participate in the emission of the same molecule — we show that the asymptotic wave dynamics of diffusive relays are governed by scaling laws. We provide a simple methodology for determining system dimensionality, and show that dimensionality dramatically affects these scaling laws. For example, cells confined to two (or one) dimensions with signaling molecule diffusion in three (or two) dimensions give rise to a diffusive wave whose speed has no dependence on *D*: a wave driven by diffusion whose speed does not depend on the rate of diffusion. In contrast to the dramatic effect of system dimensionality, these scaling laws are insensitive to biological details, including the functional form of cellular activation — the dependence of signaling molecule emission rate on the local concentration.

The formalism we use is general enough to account for the effects of signaling molecule decay, pulsed emission, chemotaxis, and the discreteness of cells, and also provides an intuitive rubric for determining under what conditions such effects alter wave propagation. Indeed, in previous seminal work, Kessler and Levine [11] have used this formalism to understand diffusive wave signaling in *Dictyostelium discoideum* with diffusion and cells in two dimensions. There, repeated pulsed emission and signaling molecule decay results in spiral signaling waves whose wave speed obeys our scaling laws only for long pulses or slow decay.

In addition to studying asymptotic wave dynamics, we systematically examine under what conditions a group of cells can trigger the formation of a diffusive wave. Here again, our results provide predictive relationships between biophysical inputs and the resulting dynamics, which are at once dramatically affected by dimensionality and largely insensitive to the details of activation.

Finally, we show that neutrophil swarming experiments [4] display dynamics consistent with our model. In this context, our results elucidate a potential design principle of diffusive relays: they create large concentration gradients. Whereas simple diffusion of a signaling molecule from a central source creates a shallow concentration profile that falls off like exp(−*r*^2^/4*Dt*), relays give rise to steep concentration profiles with gradients that quickly propagate outward and decay only modestly inside the wave front. As such, for cells like neutrophils – which use a small molecule, leukotriene B4 (LTB4), as an intercellular signaling molecule and chemoattractant (4, 18, 19) – relays provide a method for cells to collectively generate large chemical gradients that can guide directional migration. These concentration profiles, which are the result of continuous emission of a signaling molecule, are unlike those generated by repeated pulsed emission – e.g., in *Dictyostelium discoideum* [10, 11] – which create pulse trains of chemotactic cues or the flat profiles generated by relays with continuous emission and decay.

## MODEL CONSTRUCTION

We begin by considering a static group of cells uniformly distributed in two dimensions – e.g., atop a solid surface – and described by an area density *ρ* (Fig. 1A). We assume a cell at position **r** senses the local concentration of a signaling molecule, *c*(**r**, *t*), and participates in the emission at a constant rate *a* when *c*(**r**, *t*) exceeds a threshold *C*_th_. This process is depicted in Fig. 1A. Once secreted into the extracellular medium, the signaling molecules diffuse with diffusivity *D*. Treating the cells and signaling molecule in the continuum limit – we discuss the validity of doing so below with explicit details in the SI Appendix – gives rise to a single equation that governs the time evolution of *c*(**r**, *t*):

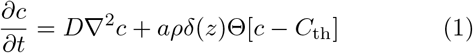

where Θ[.] is the Heaviside step function and the Dirac Delta function *δ*(*z*) accounts for the fact that the cells are confined to the plane. We discuss the validity of the latter – as well as the regime in which one can ignore signaling molecule decay – below.

**FIG. 1.**
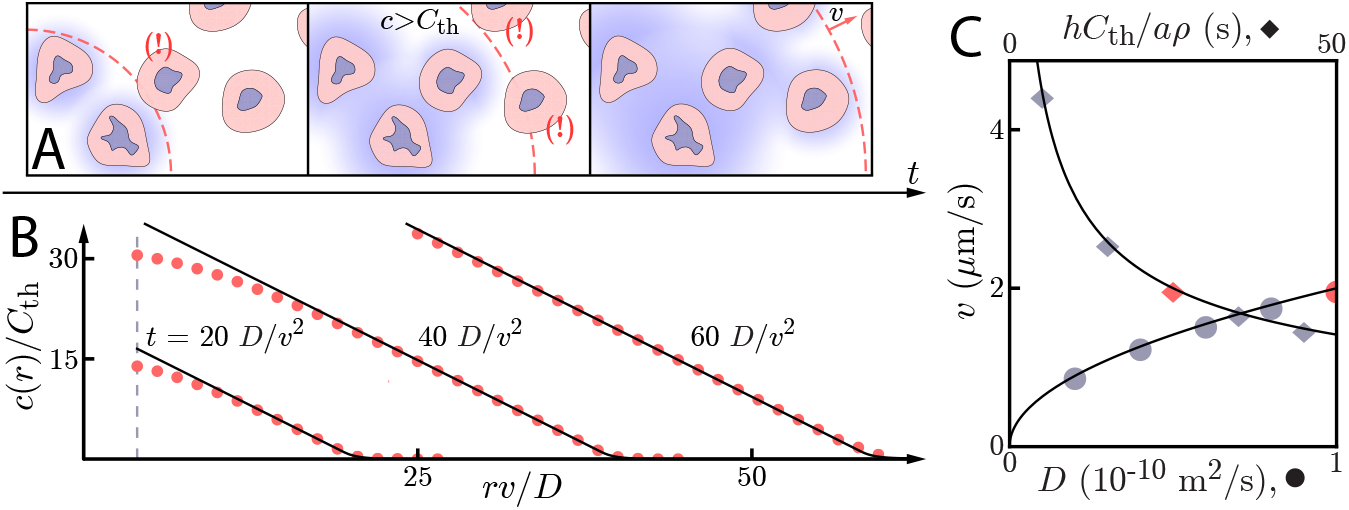
Asymptotic relay dynamics of cells in 2D with diffusion in 2D. **A:** Cartoon illustrating the diffusive relay motif. Cells (pink with purple nucleus) release a signaling molecule that diffuses (blue clouds). They do so when the local concentration exceeds a threshold, *C*_th_. This gives rise to a diffusive wave with wave speed *v*. **B:** Snapshot concentration profiles. Asymptotic theory ((5), black lines) and numerical simulation of (1) (red dots) are in good agreement and show outward-propagating waves. Here, *D* = 10^−10^ m^2^/s, *v* = 2 *μ*m/s, and *hC*_th_*/aρ* = *D/v*^2^. Numerical simulations assume that a cell colony of size *r*_*i*_ = 4*D/v* (dashed vertical line) centered at the origin starts signaling *t* = 0. **C:** Numerical wave speed as measured at *t* = 100*D/v*^2^ (markers) agrees well with theory ((4), black line) as we independently vary *D* (circles) and *hC*_th_*/aρ* (diamonds) relative to the panel **B** values (red circle and diamond).

While we at first consider cells scattered in a two-dimensional plane, one can study the signaling dynamics of cells in a one-dimensional channel or a three-dimensional environment with similar analyses. Below, we discuss the connections between the cell signaling dynamics in all of these scenarios, and all are treated in depth in the SI Appendix.

## ASYMPTOTIC WAVE DYNAMICS

Our first step in understanding diffusive signaling relays is to solve for the asymptotic dynamics of (1). Since such relays involve cells signaling their neighbors, which then signal their own neighbors, one can imagine that diffusive relays give rise to diffusive waves. We therefore make the ansatz that the concentration *c*(**r**, *t*) = *c*(*r, z, t*) can be described by an outward-traveling wave of the form 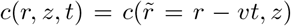 [13–15]. Here, 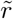 is the distance from the wave front – negative when inside the wave front, positive when beyond – and *v* is the wave speed. With 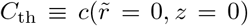 and *r* ≫ *D/v*, we take (1) and arrive at the following equation governing asymptotic behavior:

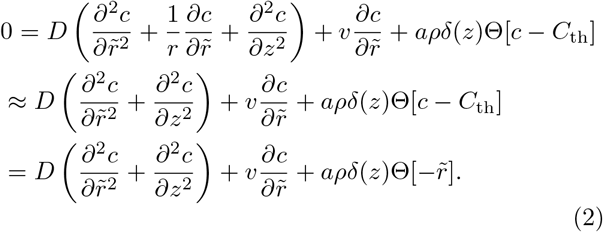

Since we consider *r* ≫ *D/v*, we may ignore the 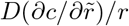 term due to the dominance of 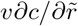. This is effectively the same as ignoring the curvature of the wave front, and has the effect of reducing our asymptotic analysis of cells in two dimensions into an asymptotic analysis of cells in one dimension [13]. The asymptotic dynamics of cells distributed in three spatial dimensions allow for a similar manipulation, an extensive discussion of which can be found in the SI Appendix.

We wish to find a solution to (2) for various diffusive – i.e., extracellular – environments. In doing so, we hope to solve for the spatial dependence of the concentration profiles 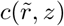 as well as a relationship that will tell us how the signaling dynamics – in this case, the wave speed *v* – depend on the biophysical system parameters like the cell density, *ρ*; the concentration threshold, *C*_th_; and the signaling molecule emission rate, *a*.

But first, we note that (2) provides two quantities of value: a natural length scale *D/v* and a natural time scale *D/v*^2^. For a small diffusing molecule with *D* ≈ 10^−10^ m^2^/s and a wave speed of *v* ≈ 1 *μ*m/s – approximately the numbers relevant for several experimental systems [1–3, 5, 10] including, as we show below, neutrophil swarming [4] – we recover *D/v* ≈ 100 *μ*m and *D/v*^2^ ≈ 100 s. We have already used the natural length scale *D/v* to derive (2) and to show that cells in 2D have the same asymptotic dynamics as cells in 1D or 3D, and we can use these scales to further justify several other approximations we have made so far. For instance, the approximation that the out-of-plane cell density can be described by *δ*(*z*) is valid when the cell thickness *H* ≪ *D/v*; similarly, decay of the signaling molecule can be neglected for a decay rate γ ≪ (*D/v*^2^)^−1^ while pulsed emission gives rise to the same asymptotic wave speed if the width of the pulse *τ* satisfies *τ* ≫ *D/v*^2^. Finally, we note that the use of (1) as a starting point is justified when the mean distance *d* between neighboring cells satisfies *dv/*4*D* ≪ 1. A thorough, mathematical discussion of all of the above, including a demonstration of why *D/v* and *D/v*^2^ are the appropriate scales, is presented in the SI Appendix. We discuss these limits below in the context of neutrophil swarming.

When the extracellular medium thickness *h* ≪ *D/v*, diffusion of the signaling molecule is effectively two-dimensional as we can take *∂*^2^*c/∂z*^2^ → 0 and *δ*(*z*) → 1*/h*. In this limit, (2) becomes

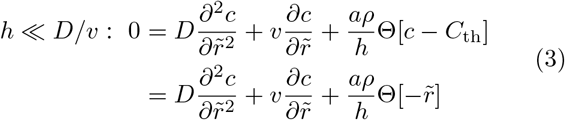

which we can solve to find the asymptotic dynamics of cells in 2D (1D, 3D) with effective signaling molecule diffusion in 2D (1D, 3D) – the thin extracellular medium limit (Fig. 1). This is the long-pulse, long-decay time limit of the model constructed by Kessler and Levine [11], and is similar to the model considered by Meyer [20]. Adding signaling molecule decay to (3) would yield a model first considered by McKean [21] in the context of nerve impulse propagation.

Before solving (3) exactly, we make two crucial observations. First, because the source (furthest right) term in (3) is proportional to *aρ/h*, all concentrations in the problem, including *C*_th_, are proportional to *aρ/h*. This gives us a single parameter to describe threshold concentration, *hC*_th_/*aρ*, which has units of time (s in SI units). The only other dimensionful parameter in the problem besides *v* – which we want to calculate – is the diffusion constant, *D*, which has units of length squared divided by time (m^2^/s in SI units). Thus, the only combination of these two parameters that will give a speed (SI units of m/s) is (*aρD/hC*_th_)^1/2^. By this simple dimensional analysis argument, the wave speed *v can only* be *v* = *α*(*aρD/hC*_th_)^1/2^ for some constant *α*. Moreover, by the same reasoning, *any* activation function – a Heaviside step function, any Hill function, a hyperbolic tangent, or anything else – that is parameterized by a single concentration *C*_th_ and emission rate *a* must give the same scalings if it has a traveling wave solution. In the SI Appendix, we numerically show that *n* ≥ 1 Hill function activation indeed gives traveling wave solutions, and that *α* ≈ 1 for *n* ≥ 2. (Also in the SI Appendix: we discuss the formal connection between Fisher’s equation [14] and (3) with Hill function activation.) The following results are therefore robust to the precise biological details of activation.

A full solution of (3), found by solving for 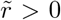 and 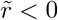 and matching boundary conditions at 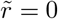, indeed reveals that

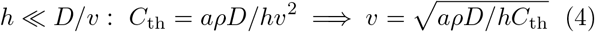

while

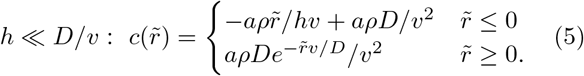

The concentration of signaling molecule thus grows linearly in the distance inside the wave front and decays exponentially in the distance beyond the wave front. We compare numerical simulations of (1) with the above asymptotic formulae for wave speeds and concentration profiles in Fig. 1B/C. For *r* ≫ *D/v*, the asymptotic formulae describe well both the concentration profile and the wave speed.

The wave speed relationship given in (4) is analogous to the Fisher-Kolmogorov wave speed [12, 14, 15] – with *hC*_th_*/aρ* replacing the doubling time as the characteristic time scale in the problem – and has been derived in beautiful previous work [12, 20] starting with Luther in 1906 [22]. In contrast to the typical 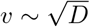 scaling seen in all these works, Vergassola *et al.* have shown that a time-dependent source term can yield an unconventional scaling of *v* ~ *D*^3/4^. Similarly, as we will now show, the dimensionality of the system can also have a dramatic effect on wave speed scaling laws.

Having solved for the asymptotic dynamics of a thin extracellular medium, we now solve for the dynamics in a thick extracellular medium for which *h* ≫ *D/v*. Here, the signaling molecules can diffuse out of plane (Fig. 2A). Because the cells sit atop a solid boundary, signaling molecules can only diffuse in the upper half of the plane and the source term in (2) acquires a factor of two to account for this boundary condition:

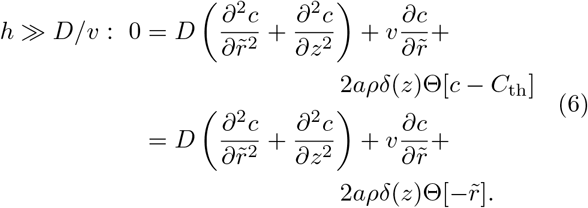

**FIG. 2.**
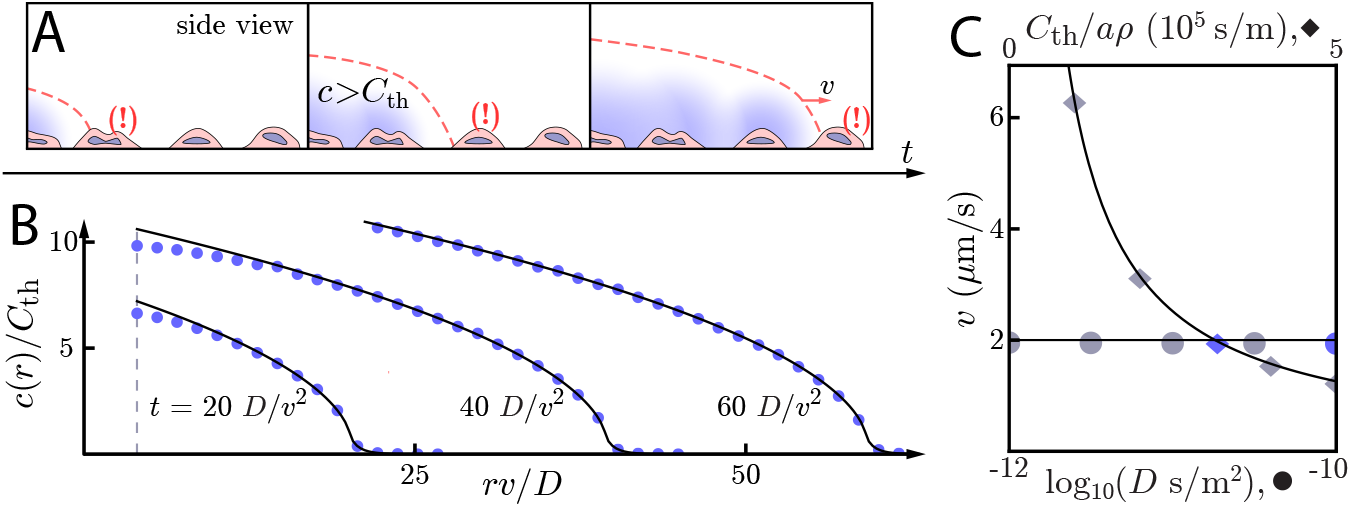
Asymptotic relay dynamics with cells in 2D and diffusion in 3D. **A:** Cartoon of cells (pink with purple nucleus) performing a diffusive relay in which signaling molecules (blue clouds) can diffuse out-of-plane. Here, such relays give rise to a diffusion-constant-independent wave speed *v*. **B:** Snapshot concentration profiles of the signaling molecule show good agreement between numerical simulation of (1) (blue dots) and asymptotic theory ((8), black lines). Here, *D* = 10^−10^ m^2^/s and *v* = 2 *μ*m/s with *C*_th_*/aρ* = 2*/πv*. The initial signaling colony is of size *r*_*i*_ = 4*D/v* (dashed vertical line). **C:** Numerical wave speed as measured at *t* = 100*D/v*^2^ (markers) agrees well with theory ((7), black line) as we independently vary *D* (circles) and *C*_th_*/aρ* (diamonds) relative to the panel **B** values (blue circle and diamond). As predicted, *v* is indeed *D*-independent in this system.

Effectively, we have cells in 2D with diffusion in 3D. We note that this case is asymptotically equivalent to cells in 1D emitting into a semi-infinite 2D environment. Thus, comparing to (5), we can see that the asymptotic dynamics are not determined by the dimension of the cell distribution or the diffusive environment, but by the *difference* in dimension between them.

Examining (6) as we did (3) reveals that every concentration in a thick extracellular medium is proportional to *aρ*. Thus, we have two dimensionful, independent parameters in (6): *C*_th_*/aρ* (SI units of s/m) and *D* (m^2^/s). The only combination of these parameters that will give a wave speed (SI units of m/s) is *aρ/C*_th_. It therefore *must* be the case that *v* = *αaρ/C*_th_ with *α* a constant – a wave driven by diffusion whose wave speed is independent of the rate of diffusion. We again stress that this is true for *any* activation function that has a traveling wave solution and can be parameterized by a single concentration *C*_th_ and a single emission rate *a*. The scaling laws governing the asymptotic dynamics are insensitive to the details of single-cell activation. As in the thin extracellular medium limit, we show in the SI Appendix that *n* ≥ 1 Hill function activation indeed gives traveling wave solutions, and that *α* ≈ 2/*π* for *n* ≥ 2.

A full solution of (6), obtained in the SI Appendix by combining a partial Fourier transform and the methods used to solve (3), yields

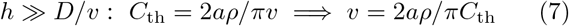

and

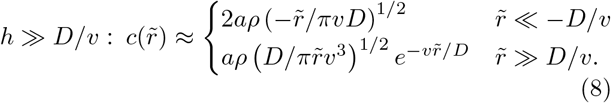

So for cells in a thick extracellular medium, the concentration grows like the square root of the distance inside the wave front and decays exponentially beyond the wave front. As with the 2D diffusive environment, we verify these relationships numerically (Fig. 2B/C). We see that the wave speed is indeed *D*-independent over two orders of magnitude in the diffusion constant.

The diffusive relay signaling motif therefore gives rise to diffusive information waves for which (4) and (7) provide predictive relationships between wave speed, threshold concentration, cell density, extracellular medium thickness, and emission rate for a variety of system dimensionalities. Similarly, (5) and (8) provide quantitative functional predictions of the concentration profiles generated by diffusive relays. By dimensional analysis, these scaling laws are insensitive to the details of activation. In the SI Appendix, we discuss the dynamics of cells in 1D with 3D diffusion as well as the properties of waves in an arbitrary extracellular medium thickness. We also explicitly account for signaling molecule decay and pulsed emission. Both have the effects of making the concentration gradient inside the wave front more gradual – or, in the case of pulsed emission with cells in 2D (or 1D) and diffusion in 3D (or 2D), creating a local concentration maximum – and of decreasing the wave speed. We emphasize that, in all cases, the asymptotic dynamics are not determined by the dimension of the diffusive or cellular environment, but by the difference in dimension between the two.

## SIGNALING WAVE INITIATION

Armed with a knowledge that diffusive relays birth diffusive waves, we now ask whether such waves are always initiated. As with the asymptotic dynamics, wave initiation depends on the system dimensionality. Here, however, the dimensionality of the diffusive environment alone determines qualitative behavior.

To demonstrate this point, we consider an “initiating colony” of cells of radius *r*_*i*_ which start signaling at *t* = 0 (Fig. 3A). As the cells expel diffusible molecules, the concentration builds up until – at the initiation time *t*_init_ – cells just outside the initiating colony (at *r*_*i*_) sense a concentration exceeding *C*_th_: *c*(*r* = *r*_*i*_, *z* = 0, *t*_init_) = *C*_th_. After *t*_init_, cells beyond *r*_*i*_ begin to participate in the diffusive relay and the diffusive wave is born.

**FIG. 3.**
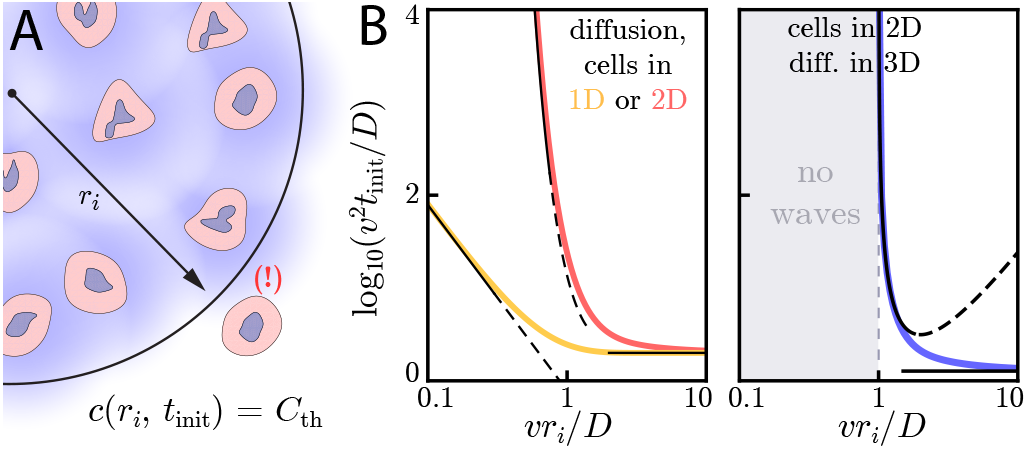
**A:** Cartoon demonstrating wave initiation. Cells within a region of size *r*_*i*_ start signaling at *t* = 0. At a later time *t* = *t*_init_, cells directly adjacent to the initial signaling region (bottom right) sense a local concentration *C*_th_. **B:** Numerics (thick colored lines) and approximate asymptotic theory ((10), thin black lines) of the initiation time’s dependence on *r*_*i*_ for cells and diffusion in 1D or 2D (left) or cells in 2D with diffusion in 3D (right). As predicted by (9), for cells and diffusion in 1D, the limit *vr*_*i*_/*D* ≪ 1 yields a dependence of 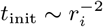 while the limit *vr*_*i*_/*D* ≫ 1 yields signaling waves that initiate at an *r*_*i*_-independent time of *t*_min,1D_ = 2*D/v*^2^. Similarly, for cells and diffusion in 2D, (10) governs the large and small *vr*_*i*_/*D* limits. In both of these cases, the wave always initiates, but the initiation time can be orders of magnitude larger than *D/v*^2^ if *r_i_* ≪ *D/v*. For cells in 2D and diffusion in 3D (right), signaling waves do not initiate for *vr*_*i*_/*D* < 1. Here again, the asymptotic theory (11) is in good agreement with numerics.

Using the Green’s function method discussed at length in the SI Appendix, we calculate how *t*_init_ varies with *D*, *v*, *r*_*i*_, and system dimensionality. This approach involves integrating Green’s functions of the diffusion equation to calculate the concentration profiles created by cells in the initiating colony. Though *v* is a composite function that depends on biophysical parameters according to (4) and (7), working with *D* and *v* allows us to invoke the natural length (*D/v*) and time (*D/v*^2^) scales mentioned above.

First, we consider cells in 1D with 1D diffusion. Here, the initiation time in the limits of small (*r*_*i*_ ≪ *D/v*) and large (*r*_*i*_ ≫ *D/v*) initiating colonies is:

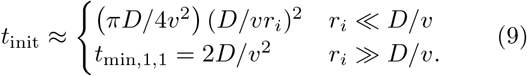

When *r*_*i*_ ≪ *D/v*, the signaling molecules quickly diffuse across and away from the initiating colony, yielding an initiation time that scales like 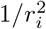. Meanwhile, for *r*_*i*_ ≫ *D/v*, the size of the initiating colony becomes irrelevant and reaches a minimum value of *t*_min,1D_, determined entirely by the characteristic time scale *D/v*^2^. The full dependence of *t*_init_ on *r*_*i*_ is pictured in Fig. 3B, where we show that the above limits are valid approximations.

Next, we consider cells in 2D with diffusion in 2D. Here, for *r*_*i*_ ≪ *D/v*, the initiation time scales harshly as

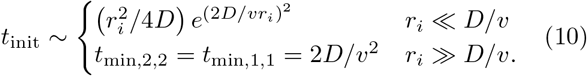

Results in the above limits are corroborated by numerical simulation in Fig. 3B, where we show initiation times for the limits above and for intermediate values of *vr_i_/D*.

Lastly, we consider cells in 2D with 3D diffusion and find that there is an initial signaling colony size of *r*_*i*_ = *πC*_th_*D/*2*aρ* = *D/v* below which the wave will not initiate. Around *r*_*i*_ = *D/v*, *t*_init_ diverges as (*vr_i_/D*)^4^(*vr_i_/D* − 1)^−2^ while for *r*_*i*_ ≫ *D/v*, *t*_init_ again plateaus at a constant value *t*_min,3D_ that only depends on the characteristic time scale *D/v*^2^:

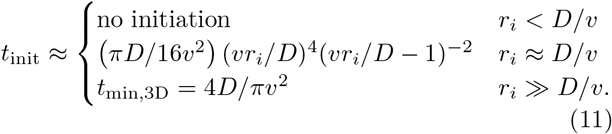

These analytic expressions agree well with numerical simulation, as seen in Fig. 3B. For cells in 3D with diffusion in 3D, there is a minimum initiating colony size of 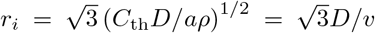. This is due to the fact that continuous sources in three-dimensional diffusive environments create steady-state concentration profiles as *t*→∞ while continuous sources in one- and two-dimensional diffusive environments do not [23]. This fact is a consequence of Polya’s Theorem, which states that that random walks in three dimensions (unlike those in one- and two-dimensions) are non-recurrent.

The critical initiating colony size for a 3D environment is reminiscent of elegant work on range expansions [13, 16]. There, the effects of diffusive migration and population growth compete with each other, and a critical mass is needed to initiate the spatial advance of a particular genotype. Here, the dimensionality of the 3D system is such that a signaling wave that will always initiate in one- and two-dimensional environments now requires a critical initial colony size.

Because the signaling wave always initiates in one- and two-dimensional environments, it can in principle be initiated by a single cell. As random activation of a single cell can initiate a signaling wave that fixes the entire population to maximal activation, these signaling dynamics have typically been thought of as unstable [24]. Yet, even in one- and two-dimensional environments, the initiation time for colonies smaller than *D/v* can be many orders of magnitude larger than the characteristic time scale of *D/v*^2^ (Fig. 3B). Thus, even though this signaling modality is technically unstable, it is robust against stochastic activation of a small number of cells over very long time scales.

As with the asymptotic wave dynamics, our conclusions are qualitatively unchanged when the cell activation is governed by a *n* > 1 Hill function instead of a Heaviside function. We catalog the initiation dynamics for all Hill functions in the SI Appendix, and have summarized the initiation and asymptotic wave dynamics for Heaviside activation in Table I.

**TABLE I.** Summary of asymptotic and initiation dynamics with Heaviside activation. One-dimensional diffusive environments are assumed to be narrow channels of width *h* in each direction perpendicular to the channel length. The cell density *ρ* has SI units 1/m for cells in 1D, 1/m^2^ for cells in 2D, and 1/m^3^ for cells in 3D. When the diffusive and cell dimensions do not match, the environment is assumed to be semi-infinite.

| | $v$ | $t_{\text{init}, \frac{vr_i}{D} \ll 1}$ | $t_{\text{init}, \frac{vr_i}{D} \gg 1}$ |
| --- | --- | --- | --- |
| cells in 1D, diff. in 1D | $\left(\frac{a\rho D}{h^2 C_{\text{th}}}\right)^{1/2}$ | $\sim \left(\frac{D}{vr_i}\right)^2$ | $2D/v^2$ |
| cells in 1D, diff. in 2D | $\frac{2a\rho}{\pi h C_{\text{th}}}$ | $\sim \exp\left(\frac{2D}{vr_i}\right)^2$ | $4D/\pi v^2$ |
| cells in 2D, diff. in 2D | $\left(\frac{a\rho D}{h C_{\text{th}}}\right)^{1/2}$ | $\sim \exp\left(\frac{2D}{vr_i}\right)^2$ | $2D/v^2$ |
| cells in 2D, diff. in 3D | $\frac{2a\rho}{\pi v}$ | no waves | $4D/\pi v^2$ |
| cells in 3D, diff. in 3D | $\left(\frac{a\rho D}{C_{\text{th}}}\right)^{1/2}$ | no waves | $2D/v^2$ |

## APPLICATION TO NEUTROPHIL SWARMING AND GRADIENT GENERATION

With a firm understanding of the diffusive wave and initiation dynamics, we now turn our sights on understanding a specific model system: neutrophil swarming. In beautiful work across several organisms [4, 25, 26], experimentalists have observed striking behavior: an acute injury or infection can elicit rapid, highly directive motion of neutrophils – the most prevalent white blood cells – towards the site of the injury or infection. These experiments have demonstrated that a lipid small molecule called leukotriene B4 (LTB4) – along with many larger, slower-diffusing proteins [4] – governs the long-range recruitment of swarming neutrophils [4, 25–27]. (Reategui *et al.* [4] have noted the presence of several other pro- and anti-inflammatory lipid small molecules during swarming, though their precise roles are less clear.) LTB4 serves to activate the neutrophils and also acts as a chemoattractant [27]; when receptors for LTB4 are blocked, swarming behavior is significantly impaired [4, 25]. The release of LTB4 has been thought to work as a relay, though the precise mechanistic details of this relay remain unclear [28, 29].

*In vitro* experiments performed with human neutrophils are particularly relevant given the results discussed so far. In these experiments, human-derived neutrophils are injected into a chamber, then settle onto the surface of a glass slide, resulting in a uniform sprinkling of cells in 2D. Also on the glass slide are circular “targets” (of size *r*_*i*_) coated in zymosan, a fungal surface protein that elicits a swarming response [4]. Some cells land on or near the target, giving an initial condition as in Fig. 3A. These cells begin signaling their neighbors, which in turn migrate towards the target (Fig. 4A). (*In vivo* experiments by Lammermann *et al.* have probed dynamics of neutrophils in complex 3D tissue environments [25].)

**FIG. 4.**
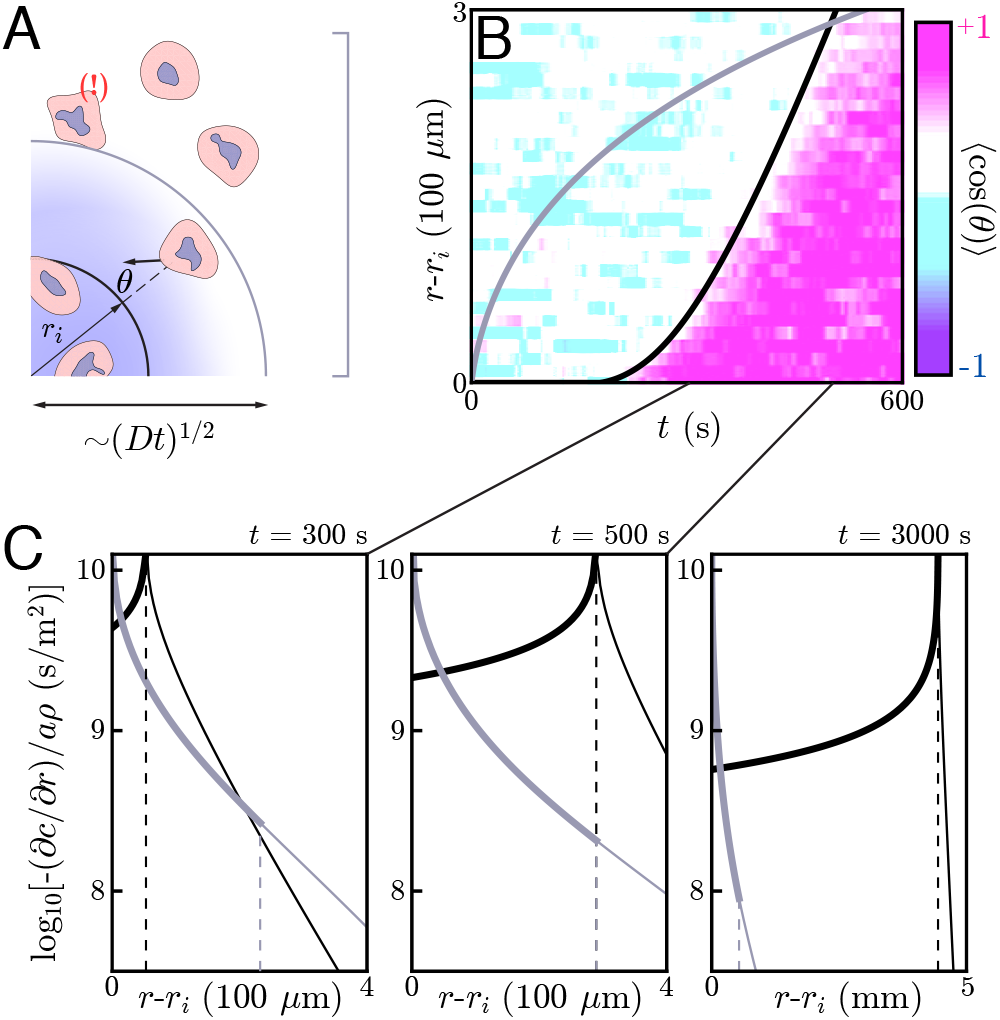
**A:** Cartoon of the simple diffusion model. Here, cells on the target (within *r_i_*) signal distant neighbors by continuously emitting a single signaling molecule. If the neighboring cells have a chemotactic response, they migrate towards the target with some noise – i.e. some non-zero angle *θ* with respect to the target. Otherwise, they move around with no sustained directionality. **B:** Experimental data (color plot) reproduced from Reategui *et al.* [4] showing the information wave front in neutrophil swarming experiments. By tracking the neutrophils in space and time, they observe highly directed motion of the neutrophils towards the target (pink) starting around *t* = 200s. There is a clear boundary in space and time – the information wave front – between the regions where cells migrate towards the target (pink) and jostle around with no particular direction (white and light blue). While a relay theory (black line) is consistent with the convex shape of the information wave front, simple diffusive signaling by only the cells on the target (grey line) is not. The diffusion constants for both models is *D* = 1.25 × 10^−10^ m^2^/s. The threshold concentrations for the relay and simple diffusion models are *C*_th_*/aρ* ≈ 3.66 × 10^5^ s/m and 2.91 × 10^4^ s/m, respectively. The parameters for the relay model are chosen to fit the wave front by eye while the simple diffusion model parameters are chosen to give the same signaling distance at *t* = 500 s. **C:** Gradients created by signaling relays (black) and simple diffusion (grey) models in panel B. The dashed vertical lines indicate the location of the information wave front. As time increases from left to right, the relay signaling motif gives an information wave that signals cells faster than simple diffusion in the long time limit. Cells within the wave front (solid lines; dashed lines indicate regions beyond the wave fronts) experience significantly larger gradients when the cells utilize a relay, which may facilitate efficient chemotaxis.

By tracking individual cells in time, one can deduce their migratory direction as a function of time. A typical metric for quantifying the directionality a cell’s migration is the chemotactic index – the cosine of the angle *θ* between a cell’s motion and the direction of the target (Fig. 4A). One can average over the cells at a given distance *r* and time *t* to construct a plot of the average directionality ⟨cos *θ*⟩ in space and time. As pictured in Fig. 4B, such a plot reveals a clear divide in space and time between cells that are highly directed towards the target (pink) and those without any particular directionality (white and light blue). We refer to the boundary of this divide as an information wave front – cells that lie underneath the curve have received the signal and begun chemotaxing towards the target while those above the curve have not.

Interestingly, the information wave front is convex with respect to the origin – a dramatic departure from what simple diffusive signaling by cells on the target would yield (Fig. 4A/B). We therefore posit that the cells may be participating in a relay in which they emit LTB4 in response to the same and check to see if this is consistent with the observed information wave front.

To do so, we perform a numerical simulation of (1) with an additional term to account for the signaling of cells that land on the target. For this analysis, we assume a circular target of radius *r*_*i*_ ≈ 100 *μ*m, though the targets fabricated by Reategui *et al.* are smaller, oblong objects. Here, the diffusive environment is effectively three dimensional and the cells are close enough to allow for the use of a continuum model like (1) (see below). Our model assumes switch-like activation of neutrophils, which we associate with the onset of directed chemotaxis. While we ignore the inward migration of cells in this analysis, we show that it has a negligible effect on the information wave propagation in the SI Appendix. Thus, as mentioned above, (1) effectively has two parameters: *C*_th_*/aρ* and *D*. Fitting these two parameters to the observed information wave front gives *C*_th_*/aρ* ≈ 3.67 × 10^5^ s/m and *D* ≈ 1.25 × 10^−10^ m^2^/s, the latter of which is consistent with the diffusion constant of a small molecule like LTB4. This implies a wave speed of *v* ≈ 1.7*μ*m/s. Thus, we are validated in using a continuum model with a thick extracellular medium, as for this experiment the extracellular medium thickness *h* = 2 mm ≫ *D/v* and the mean distance between neutrophils, *d* = 50 *μ*m, satisfies *vd/*4*D* ≈ 0.17 ≪ 1. The cell thickness *H* ≈ 10 *μ*m indeed satisfies *H* ≪ *D/v*, meaning the use of the delta function to describe the cell distribution is valid. Finally, as LTB4 has a lifetime 1/γ of many minutes [30] and *D/v*^2^ ≈ 40 s ≪ 1/γ, we can indeed ignore signaling molecule decay. These fit parameters give a curve that matches the transient dynamics over the field of view of the experiment (Fig. 4). Thus, our relay model gives dynamics that are consistent with the dynamics of neutrophil swarming experiments – namely, the observed convex shape of the information wave front. Larger field-of-view and longer time course experiments with varying cell densities and larger targets will provide a deeper mechanistic understanding of such relays, while also testing the scaling predictions of (4) and (7).

Finally, we seek an answer to the question of *why* neutrophils might employ such signaling relays. As we have shown above, relays lead to “fast” communication, in the sense that they give rise to diffusive waves which travel a distance *vt* in a time *t*, compared to the 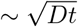 distance of simple diffusion. However, there is another potential design principle underlying diffusive relays: they create strong gradients that may help cells chemotax effectively, as we will now demonstrate.

To get an idea of the gradients we are working with, we compare those generated by a relay – calculated by solving (2) and approximated in (8) – to a comparable simple diffusion model, such as that pictured in 4B. (In the SI Appendix, we present the same comparison for a thin extracellular medium.) As is well-known, a burst-like emission of a diffusible molecule creates shallow, Gaussian concentration profiles away from the source; the same is true for continuous emission of a fixed source (see SI Appendix). Thus, the gradients that individual cells or small colonies of cells can create through simple diffusive signaling are orders of magnitude shallower than the collective gradients generated by relays (Fig. 4C). This hints that neutrophils may use relays not solely for their improved signaling speed, but also for the strong resulting chemotactic gradients.

## DISCUSSION

In this work, we have shown how simple cell signaling relays can give rise to diffusive waves whose properties are robust to many underlying details. Our work especially highlights the importance of the dimensionality of the extracellular medium, as seemingly innocent changes to the environment can have large effects on the resulting diffusive waves. The strong effect of system dimensionality is reminiscent of previous work on diffusive dynamics, which showed how dimensionality can effect Turing pattern instabilities [31].

Though we have characterized the asymptotic dynamics, initiation, and potential design principles of these waves in several scenarios, many interesting problems remain as yet unsolved. First, as noted by Lammermann and colleagues [28, 29], it is unclear how the complexities of *in vivo* extracellular environments affect these results, particularly in the context of neutrophil swarming. Ambient flow (e.g., in blood vessels), constrictions, and complex diffusive environments may lead to dynamics of biological relevance beyond those discussed here. Additionally, it would be interesting to study how different models of chemoreception and cellular uptake – topics of theoretical [19] and experimental [32–34] relevance – affect our conclusions.

It would also be interesting to leverage the design principles we have discussed for engineering synthetic relays, a field with a rich history [5, 35–37]. To that end, our results provide a general framework for determining how system dimensionality, diffusion constants, activation functions, cell density, etc. affect cell signaling and wave initiation. Experimental work on this problem and others would provide tests of our many quantitative predictions.

## Acknowledgements

We thank Richard Murray, James Parkin, Justin Bois, Mikhail Shapiro, David Nelson, Taekjip Ha, and Jenny Sheng for helpful discussions and Felice Frankel for help with figure preparation. P.B.D. is supported by the Paul M. Young Fellowship through the Fannie and John Hertz Foundation, the NSF through MRSEC DMR 14-20570, and the Kavli Foundation.

