## Supplementary material for "Dynamics of diffusive cell signaling relays": SI

#### 1 Model set-up

Before doing any math, let's set up the scenarios we intend to study. We consider a continuum of cells described by a cell density $\rho$. These cells emit one type of signaling molecule at a rate $a$ when the local concentration of the signaling molecule is above a certain threshold, $C_{\mathrm{th}}$. The molecules diffuse in the extracellular medium with diffusion constant $D$. The concentration of the signaling molecule is described by the variable $c$, which is a function of both space and time: $c = c(\mathbf{r}, t)$. In general, then, we have

$$\frac{\partial c}{\partial t} = D\nabla^2 c + a\rho\Theta[c - C_{\mathrm{th}}] \tag{S1}$$

with $\Theta[.]$ the Heaviside step function. We study this model and variants going forward.

#### 2 Asymptotic wave ansatz

We start out by seeking to understand what dynamical properties a system described by (S1) has at large times. To study these dynamics, we need an inspired guess for what the dynamics will look like. We imagine that when a small volume of cells starts signaling its neighbors – and those neighbors start signaling their neighbors – that a reasonable guess for the dynamics of such a signaling relay is an outward propagating wave with speed $v$. We define $r$ as the distance from the center of the outward propagating wave. At long times for a uniform cell density, the shape information wave front will obey radial symmetry and our ansatz becomes $c(\mathbf{r}, t) = c(\mathbf{r} - vt\hat{r})$ with $\hat{r}$ the unit vector pointing from the origin to the wave front.

With this guess, we can define a new coordinate $\tilde{r} = r - vt$ (we call this $\tilde{x} = x - vt$ for cells in one dimension) which defines the distance to the wave front. Note that $\tilde{r} < 0$ means we are inside the wave front while $\tilde{r} > 0$ means we are beyond it. With these definitions, $\partial c/\partial\tilde{r} = \partial c/\partial r$ and $\partial c/\partial t = -v\ \partial c/\partial\tilde{r}$. For cells in 1D, we consider $y$ and $z$ to be dimensions perpendicular to the line of cells with the density described by $\rho\delta(y)\delta(z)$ with $\rho$ measured in cells per unit length; for cells in 2D, we consider $z$ to be the out-of-plane dimension and the density to be described by $\rho\delta(z)$ with $\rho$ measured in cells per unit area; for cells in 3D, $\rho$ is measured in cells per unit volume. Assuming azimuthal symmetry in 2D and radial symmetry in 3D, we arrive at

$$\text{cells in 1D: } 0 = D\left(\frac{\partial^2 c}{\partial\tilde{x}^2} + \frac{\partial^2 c}{\partial y^2} + \frac{\partial^2 c}{\partial z^2}\right) + v\frac{\partial c}{\partial\tilde{x}} + a\rho\ \delta(y)\delta(z)\Theta[c - C_{\mathrm{th}}] \tag{S2a}$$

$$\text{cells in 2D: } 0 = D\left(\frac{\partial^2 c}{\partial\tilde{r}^2} + \frac{1}{r}\frac{\partial c}{\partial\tilde{r}} + \frac{\partial^2 c}{\partial z^2}\right) + v\frac{\partial c}{\partial\tilde{r}} + a\rho\ \delta(z)\Theta[c - C_{\mathrm{th}}] \tag{S2b}$$

$$\text{cells in 3D: } 0 = D\left(\frac{\partial^2 c}{\partial\tilde{r}^2} + \frac{2}{r}\frac{\partial c}{\partial\tilde{r}}\right) + v\frac{\partial c}{\partial\tilde{r}} + a\rho\ \Theta[c - C_{\mathrm{th}}] \tag{S2c}$$

These equations can be simplified once more by noting that we are considering asymptotic – i.e., large $r$ – dynamics. Thus, as long as $v \gg D/r$, we can say that $v\partial c/\partial\tilde{r}$ dominates terms like $D\left(\partial c/\partial\tilde{r}\right)/r$ and we can

*

ignore the latter [1]. By construction, our ansatz says that $c(\tilde{r}=0) \equiv C_{\rm th}$, meaning that $\Theta[c - C_{\rm th}] = \Theta[-\tilde{r}]$. This gives simplified equations according to:

$$\text{cells in 1D: } 0 = D\left(\frac{\partial^2 c}{\partial \tilde{x}^2} + \frac{\partial^2 c}{\partial y^2} + \frac{\partial^2 c}{\partial z^2}\right) + v\frac{\partial c}{\partial \tilde{x}} + a\rho\ \delta(y)\delta(z)\Theta[-\tilde{x}] \tag{S3a}$$

$$\text{cells in 2D: } 0 = D\left(\frac{\partial^2 c}{\partial \tilde{r}^2} + \frac{\partial^2 c}{\partial z^2}\right) + v\frac{\partial c}{\partial \tilde{r}} + a\rho\ \delta(z)\Theta[-\tilde{r}] \tag{S3b}$$

$$\text{cells in 3D: } 0 = D\frac{\partial^2 c}{\partial \tilde{r}^2} + v\frac{\partial c}{\partial \tilde{r}} + a\rho\ \Theta[-\tilde{r}]. \tag{S3c}$$

These equations provide both a natural length scale, $D/v$, and a natural timescale, $D/v^2$. One can see that these are the relevant time and length scales for our problem by non-dimensionalizing, e.g., (S3c) to get

$$\text{cells in 3D: } 0 = \frac{\partial^2 (cv^2/a\rho D)}{\partial (v\tilde{r}/D)^2} + \frac{\partial (cv^2/a\rho D)}{\partial (v\tilde{r}/D)} + \Theta[-v\tilde{r}/D]. \tag{S4}$$

Thus, every length scale in the problem is normalized by $D/v$; because of our traveling wave ansatz, every length scale can be converted to a time scale by dividing by $v$, giving $D/v^2$ as the natural timescale. The natural length scale is useful, e.g., for understanding what it means for cells to be in "one dimension" or for diffusion to be in "two dimensions". If the cells are organized in a line (or on a plane) such that their average deviation from the line (or distance from the plane) is $d \ll D/v$, then they are effectively in one (or two) dimensions and (S3a) (or (S3b)) holds. If cells are constricted to a narrow channel of width $h \ll D/v$ (or an extracellular medium of thickness $h \ll D/v$), then diffusion is effectively one (or two) dimensional. For instance, for cells confined in a very narrow one-dimensional channel of width $h \ll D/v$, we can simplify (S3a) because $\partial^2 c/\partial y^2 = \partial^2 c/\partial z^2 = 0$ and $\delta(y)\delta(z) \to 1/h^2$. The resulting equation is the exact same as for cells in 3D but with a source term proportional to $a\rho/h^2$:

$$\text{cells in 1D, diffusion in 1D: } 0 = D\frac{\partial^2 c}{\partial \tilde{x}^2} + v\frac{\partial c}{\partial \tilde{x}} + \frac{a\rho}{h^2}\Theta[-\tilde{x}]. \tag{S5}$$

For cells in 2D with an extracellular medium of thickness $h \ll D/v$, diffusion is effectively two-dimension and, by the same logic that produced (S5), the asymptotic governing equation is:

$$\text{cells in 2D, diffusion in 2D: } 0 = D\frac{\partial^2 c}{\partial \tilde{r}^2} + v\frac{\partial c}{\partial \tilde{r}} + \frac{a\rho}{h}\Theta[-\tilde{r}]. \tag{S6}$$

As (S4), (S5), and (S6) are all the same, the dynamics of cells in 1D with diffusion in 1D are the same as those of cells in 2D with diffusion in 2D or those in 3D with diffusion in 3D.

One can, however, arrive at a different governing equation by considering cells in 2D (e.g., cells sitting on a plane) with a thick extracellular medium of thickness $h \gg D/v$. In this case, diffusion effectively takes place in three dimensions and

$$\text{cells in 2D, diffusion in 3D: } 0 = D\left(\frac{\partial^2 c}{\partial \tilde{r}^2} + \frac{\partial^2 c}{\partial z^2}\right) + v\frac{\partial c}{\partial \tilde{r}} + a\rho\delta(z)\Theta[-\tilde{r}]. \tag{S7}$$

which, by the same logic above, is functionally equivalent to the governing equation for cells in 1D with diffusion in 2D. We can therefore see that it is not the dimensionality of the cell distribution or the diffusive environment that determines the asymptotic dynamics, but rather the difference in dimension between the two.

Going forward, we will think of cells in two dimensions, as we have done in the main text. This will allow us to interpolate between an effectively two-dimensional diffusive environment (the thin extracellular medium limit) and an effectively three-dimensional environment (the thick extracellular medium limit).

#### Cells in 2D, diffusion in 2D: the thin extracellular medium limit

For an extracellular medium of thickness $h \ll D/v$, diffusion effectively takes place in two dimensions as argued in the previous section. The signaling molecule concentration has no $z$-dependence and concentrations get normalized by $h$. Here,

$$0 = D\frac{\partial^2 c}{\partial \tilde{r}^2} + v\frac{\partial c}{\partial \tilde{r}} + \frac{a\rho}{h}\Theta[-\tilde{r}] \tag{S8}$$

is our asymptotic governing equation.

For both $\tilde{r} < 0$ and $\tilde{r} > 0$, (S8) reduces to two straightforward-to-solve linear ODEs. With $b_i$ as constants that we will determine momentarily,

$$\tilde{r} < 0: \; 0 = D\frac{\partial^2 c}{\partial \tilde{r}^2} + v\frac{\partial c}{\partial \tilde{r}} + a\rho/h \implies c(\tilde{r} < 0) = b_2 e^{-v\tilde{r}/D} + b_3 - a\rho\tilde{r}/hv \tag{S9a}$$

$$\tilde{r} > 0: \; 0 = D\frac{\partial^2 c}{\partial \tilde{r}^2} + v\frac{\partial c}{\partial \tilde{r}} \implies c(\tilde{r} > 0) = b_0 e^{-v\tilde{r}/D} + b_1. \tag{S9b}$$

We can solve for the $b_i$ by applying boundary conditions. First, we demand $c \to 0$ as $\tilde{r} \to \infty$ and that the concentration profile only blow up linearly as $\tilde{r} \to -\infty$. Physically, these demands are justified as follows: the concentration as $\tilde{r} \to \infty$ has to go to zero because there are no cells emitting in that region and it is far from the wave front; the concentration as $\tilde{r} \to -\infty$ can grow at most linearly because the cells a distance $-\tilde{r}$ from the wave front have only been emitting for a time $-\tilde{r}/v$. Combining these asymptotic boundary conditions with the demand that $c(\tilde{r})$ be continuous and have a continuous first derivative at $\tilde{r} = 0$ allows us to stitch together the solutions in (S9a) and (S9b) to yield:

$$c(\tilde{r} < 0) = a\rho D/hv^2 - a\rho\tilde{r}/hv \tag{S10a}$$

$$c(\tilde{r} > 0) = a\rho D e^{-v\tilde{r}/D}/hv^2. \tag{S10b}$$

We show this concentration profile in the left panel of Fig. 1A. From the above, we infer that

$$C_{\text{th}} = c(0) = a\rho D/hv^2 \implies v = \sqrt{a\rho D/hC_{\text{th}}}. \tag{S11}$$

Thus, we have an explicit formula relating wave speed, emission rate, cell density, diffusion constant, extracellular medium thickness, and threshold concentration. We can also see that the concentration profile beyond the wave front is exponential, not Gaussian as for simple diffusion. The concentration inside the wave front grows linearly as the distance from the wave front.

#### Cells in 2D, diffusion in 3D: the thick extracellular medium limit

Next, we consider (S3b) in the limit that the extracellular medium $h \gg D/v$. In this limit, we effectively have cells in 2D with diffusion in 3D. With cells sitting on a substrate, signaling molecules can only diffusive in the upper half of the plane, and we have a semi-infinite environment which accounts for an extra factor of 2 in the emission term, yielding:

$$0 = D\left(\frac{\partial^2 c}{\partial \tilde{r}^2} + \frac{\partial^2 c}{\partial z^2}\right) + v\frac{\partial c}{\partial \tilde{r}} + 2a\rho\, \delta(z)\Theta[-\tilde{r}]. \tag{S12}$$

Instead of working directly with $\delta(z)$, we consider the cells to be of a thickness $H$ such that $2\delta(z) \to \frac{1}{H}\sqrt{\frac{2}{\pi}}\exp(-z^2/2H^2)$ and

$$0 = D\left(\frac{\partial^2 c}{\partial \tilde{r}^2} + \frac{\partial^2 c}{\partial z^2}\right) + v\frac{\partial c}{\partial \tilde{r}} + \frac{a\rho}{H}\sqrt{\frac{2}{\pi}}e^{-z^2/2H^2}\Theta[-\tilde{r}] \tag{S13}$$

Next, we take a partial Fourier transform of (S12) with $k$ and $C(\tilde{r},k)$ the Fourier partners of $z$ and $c(\tilde{r},z)$, respectively[1]. This gives:

$$0 = D\left(\frac{\partial^2 C}{\partial \tilde{r}^2} - k^2 C\right) + v\frac{\partial C}{\partial \tilde{r}} + \sqrt{\frac{2}{\pi}} e^{-H^2k^2/2} a\rho\ \Theta[-\tilde{r}] \tag{S14}$$

which is another pair of piecewise, straightforward-to-solve, linear ODEs. Solving with the same boundary conditions that yielded (S10a) and (S10b), we arrive at

$$C(\tilde{r}<0,k) = \frac{a\rho e^{-H^2k^2/2}}{Dk^2\sqrt{2\pi}}\left(2 - \frac{\left(1+\sqrt{4D^2k^2/v^2+1}\right)\exp\left(\frac{v\tilde{r}}{2D}\left(\sqrt{4D^2k^2/v^2+1}-1\right)\right)}{\sqrt{4D^2k^2/v^2+1}}\right) \tag{S15a}$$

$$C(\tilde{r}>0,k) = \frac{a\rho e^{-H^2k^2/2}}{Dk^2\sqrt{2\pi}}\frac{\sqrt{4D^2k^2/v^2+1}-1}{\sqrt{4D^2k^2/v^2+1}}\exp\left(-\frac{v\tilde{r}}{2D}\left(\sqrt{4D^2k^2/v^2+1}+1\right)\right). \tag{S15b}$$

To find the concentrations at the cells, we can take the inverse partial Fourier transform of these expressions at $z=0$. But first, we note that the right sides of (S15a) and (S15b) have no support when $k \gg v/D$. Thus, if $Hv/D \ll 1$, the term $e^{-H^2k^2/2}$ is irrelevant for calculating the real-space concentrations and can be replaced with 1. This is equivalent to having chosen $\delta(z)$ to describe the out-of-plane cell density.

Proceeding with $e^{-H^2k^2/2} \to 1$, one can take the inverse partial Fourier transform and arrive at

$$C_{\text{th}} = 2a\rho/\pi v. \tag{S16}$$

Similarly, in the limit $|\tilde{r}| \gg D/v$,

$$c(\tilde{r} \ll -D/v, z) \approx \frac{2a\rho}{v}\sqrt{\frac{-\tilde{r}v}{\pi D}}\left(e^{vz^2/4D\tilde{r}} - \sqrt{-\frac{\pi v z^2}{4D\tilde{r}}}\operatorname{erfc}\sqrt{-\frac{vz^2}{4D\tilde{r}}}\right) \tag{S17a}$$

$$c(\tilde{r} \gg D/v, z) \approx a\rho\sqrt{\frac{D}{\pi\tilde{r}v^3}}e^{-v\tilde{r}/D}e^{-vz^2/4D\tilde{r}}. \tag{S17b}$$

The former has the same functional dependence on $z$ as the concentration a distance $z$ away from a continuously emitting point source with diffusion in 1D after a time $\tilde{r}/v$ (see Section 6). Using the above, we can find the concentration in the plane of the cells ($z=0$):

$$c(\tilde{r} \ll D/v, z=0) \approx \frac{2a\rho}{v}\sqrt{\frac{-\tilde{r}v}{\pi D}} \tag{S18a}$$

$$c(\tilde{r} \gg D/v, z=0) \approx a\rho\sqrt{\frac{D}{\pi\tilde{r}v^3}}e^{-v\tilde{r}/D}. \tag{S18b}$$

We show this concentration profile in the left panel of Fig. 1B.

#### Cells in 1D, diffusion in 3D: an artificial case

Finally, we consider a line of cells in one dimension with diffusion taking place in three dimensions. This corresponds to a somewhat artificial test case of cells in a line with mean distance from the line $d \ll D/v$ and diffusion in an environment of size $h \gg D/v$ in the dimensions perpendicular to this line of cells. Nonetheless, it is interesting because we have to include the finite size of the cells in order to get a traveling wave solution. Here,

$$0 = D\left(\frac{\partial^2 c}{\partial \tilde{x}^2} + \frac{\partial^2 c}{\partial y^2} + \frac{\partial^2 c}{\partial z^2}\right) + v\frac{\partial c}{\partial \tilde{x}} + \frac{a\rho}{\pi H^2}e^{-\left(y^2+z^2\right)/2H^2}\Theta[-\tilde{r}] \tag{S19}$$

[1]Here, we choose $C(\tilde{r},k) \equiv \frac{1}{\sqrt{2\pi}}\int_{-\infty}^{\infty} e^{ikz}c(\tilde{r},z)$

with $v$ the wave speed; $\tilde{x}$ the distance to the wave front; $y$ and $z$ the extra diffusive dimensions; and $H$ the size of the cells.

Taking partial Fourier transforms across both $y$ and $z$ gives with the Fourier transform conventions and notation used above gives

$$0 = D\left(\frac{\partial^2 C}{\partial \tilde{x}^2} - k_y^2 C - k_z^2 C\right) + v\frac{\partial C}{\partial \tilde{x}} + \frac{a\rho}{\pi} e^{-\left(k_y^2+k_z^2\right)H^2/2}\Theta[-\tilde{x}] \tag{S20}$$

which reduces to (S14) with $k^2 \to k_y^2 + k_z^2$. One can then find the concentration profiles and self-consistency relationship for $C_{\mathrm{th}}$ by inverse Fourier transforming the analogs of (S18a) and (S18b). The value of $C_{\mathrm{th}} = c(\tilde{x} = 0, y = z = 0)$ diverges as $H \to 0$.

### 3 Pulsed emission and decay

In this section, we consider pulsed emission and decay of the signaling molecule. These scenarios are relevant for signaling pathways in, e.g., *Dictyostelium* [2–4] and *E. Coli* [5], in which intracellular dynamics produce a pulse-like release of signaling molecules into the extracellular medium. Kessler and Levine [4] have previously used this machinery to construct a signaling model for Dictyostelium, including pulsed emission and signaling molecule decay. Here, we consider the effects of each independently.

We explicitly discuss only the asymptotics of cells in 2D with diffusion in 2D (equivalent to cells in 1D with diffusion in 1D or cells in 3D with diffusion in 3D, as shown previously), though we quote the results for cells in 2D with diffusion in 3D (equivalent to cells in 1D with diffusion in 2D) which are obtained using the Fourier transform machinery in Section 2. Here again, the asymptotic dynamics depend on the difference in dimensionality between the cellular and the diffusive environment.

#### Pulsed emission with cells in 2D and diffusion in 2D

Here, we consider a square pulse of length $\tau$ emitted once a cell exceeds the threshold concentration $C_{\mathrm{th}}$. In the moving frame, this pulse has length $v\tau$ – for notational simplicity here, we dispense with dimensional subscripts on the wave speed – giving rise to the pulsed emission analog of (S8):

$$0 = D\frac{\partial^2 c}{\partial \tilde{r}^2} + v\frac{\partial c}{\partial \tilde{r}} + \frac{a\rho}{h}\Theta[-\tilde{r}]\Theta[\tilde{r} + v\tau] \tag{S21}$$

which is nothing more than three piecewise linear equations, which we stitch together as before. The source term is zero when $\tilde{r} < -v\tau$ (Region I) or $\tilde{r} > 0$ (Region III) and $a\rho_0$ for $-v\tau < \tilde{r} < 0$ (Region II). We thus recover the following:

$$\text{Region I: } c(\tilde{r} < -v\tau) = b_1 + b_2 e^{-v\tilde{r}/D} \tag{S22a}$$

$$\text{Region II: } c(-v\tau < \tilde{r} < 0) = b_3 + b_4 e^{-v\tilde{r}/D} - a\rho\tilde{r}/hv \tag{S22b}$$

$$\text{Region III: } c(x > 0) = b_5 + b_6 e^{-v\tilde{r}/D} \tag{S22c}$$

Applying the same boundary conditions as with continuous emission, we arrive at

$$\text{Region I: } c(\tilde{r} < -v\tau) = a\rho\tau/h \tag{S23a}$$

$$\text{Region II: } c(-v\tau < \tilde{r} < 0) = a\rho D/hv^2 - \frac{a\rho D}{hv^2} e^{-v^2\tau/D} e^{-v\tilde{r}/D} - a\rho\tilde{r}/hv \tag{S23b}$$

$$\text{Region III: } c(\tilde{r} > 0) = \frac{a\rho D}{hv^2}(1 - e^{-v^2\tau/D})e^{-v\tilde{r}/D} \tag{S23c}$$

which tells us a few things of interest. First – as seen in the right panel of Fig. 1 A (here, $\tau = 2D/v^2 = 2C_{\mathrm{th}}/a\rho$) – the concentration profile for $\tilde{r} < -v\tau$ is flat. (As with continuous emission, pulsed emission gives the familiar exponential profile beyond the wave front.) Second, the wave speed, pulse width, cell density, emission rate, extracellular medium thickness, and threshold concentration are related through the equation

$$C_{\rm th} = a\rho D(1 - e^{-v^2\tau/D})/hv^2. \tag{S24}$$

For $\tau \gg D/v^2$, we recover the usual relationship of $C_{\rm th} = a\rho D/hv^2$. In region 2, the profile will grow linearly as before until $-v\tilde{r}/D$ becomes comparable to $v^2\tau/D$, at which point $e^{-v^2\tau/D - v\tilde{r}/D}$ becomes of order unity and the profile levels off.

To understand how the wave speed with pulsed emission, $v$, compares to the wave speed with continuous emission, $(a\rho D/hC_{\rm th})^{1/2}$, we have plotted $v/(a\rho D/hC_{\rm th})^{1/2}$ as a function of $1/\tau$ in Fig. 2A. We have normalized $\tau$ by a characteristic time $\tau_c = hC_{\rm th}/a\rho$, which is equal to $D/v^2$ for continuous emission. When $\tau < \tau_c$, the wave speed goes to zero. There is no wave-like solution for shorter pulses.

We note that a timed pulsed emission considered here is formally equivalent to cells signaling until the local concentration exceeds $c(\tilde{r} = -v\tau) = a\rho\tau/h$. This is relevant in, e.g., quorum sensing models in which the local presence of a signaling molecule can both upregulate (at relatively low concentrations) and downregulate (at relatively high concentrations) release of the same signaling molecule [5].

#### Continuous emission plus decay with cells in 2D and diffusion in 2D

At last, we characterize the effect of signaling molecule decay at rate $\gamma$ by adding a term of $-\gamma c$ to (S3b). In the thin extracellular medium limit,

$$0 = D\frac{\partial^2 c}{\partial \tilde{r}^2} + v\frac{\partial c}{\partial \tilde{r}} + \frac{a\rho}{h}\Theta[-\tilde{r}] - \gamma c. \tag{S25}$$

This is another piecewise set of linear differential equations, which we can solve as without decay to yield:

$$c(\tilde{r} < 0) = \frac{a\rho}{h\gamma} - \frac{a\rho}{2h\gamma}\left[1 + (1 + 4D\gamma/v^2)^{-1/2}\right]\ e^{\frac{\tilde{r}v}{2D}\left(\sqrt{4D\gamma/v^2+1}-1\right)} \tag{S26a}$$

$$c(\tilde{r} > 0) = \frac{a\rho}{2h\gamma}\left[1 - (1 + 4D\gamma/v^2)^{-1/2}\right]\ e^{-\frac{\tilde{r}v}{2D}\left(\sqrt{4D\gamma/v^2+1}+1\right)} \tag{S26b}$$

as the concentration profiles and

$$C_{\rm th} = \frac{a\rho}{2h\gamma}\left[1 - (1 + 4D\gamma/v^2)^{-1/2}\right] \implies v = 4D\gamma\left[\left(1 - \frac{2h\gamma C_{\rm th}}{a\rho}\right)^{-2} - 1\right]^{-1} \tag{S27}$$

as our wave speed relationship. The concentration profile is flatter than its decay-free counterpart (Fig. 1A). For $\gamma \ll v^2/D$, (S27) gives the decay-free relationship. And – as with pulsed emission – the wave speed goes to zero, this time when $\gamma \to 1/2\tau_c$ where $\tau_c = hC_{\rm th}/a\rho$ (Fig. 2B).

#### Pulsed emission with cells in 2D and diffusion in 3D

Here,

$$0 = D\left(\frac{\partial^2 C}{\partial \tilde{r}^2} - k^2 C\right) + v\frac{\partial C}{\partial \tilde{r}} + \sqrt{2/\pi}\ a\rho\ \Theta[-\tilde{r}]\Theta[v\tau + \tilde{r}] \tag{S28}$$

which we can solve to yield:

$$\text{Region I: } C(\tilde{r} < -v\tau, k) = B_1(k)e^{\frac{\tilde{r}v}{2D}\left(-1+\sqrt{1+4D^2k^2/v^2}\right)} \tag{S29a}$$

$$\text{Region II: } C(-v\tau < \tilde{r} < 0, k) = \sqrt{\frac{2}{\pi}}\frac{a\rho_0}{Dk^2} + B_2(k)e^{-\frac{\tilde{r}v}{2D}\left(1+\sqrt{1+4D^2k^2/v^2}\right)} + B_3(k)e^{\frac{\tilde{r}v}{2D}\left(-1+\sqrt{1+4D^2k^2/v^2}\right)} \tag{S29b}$$

$$\text{Region III: } C(\tilde{r} > 0, k) = B_4(k) e^{-\frac{\tilde{r}v}{2D}\left(1+\sqrt{1+4D^2k^2/v^2}\right)} \tag{S29c}$$

with $B_i(k)$ chosen such that $C$ and its first derivative are continuous:

$$B_1(k) = \frac{a\rho}{D\sqrt{2\pi}} \frac{\left(v + \sqrt{4D^2k^2 + v^2}\right)\left(e^{\frac{v^2\tau}{2D}\left(\sqrt{4D^2k^2/v^2+1}-1\right)} - 1\right)}{k^2\sqrt{4D^2k^2 + v^2}} \tag{S30a}$$

$$B_2(k) = \frac{a\rho}{D\sqrt{2\pi}} \frac{\left(v - \sqrt{4D^2k^2 + v^2}\right) e^{-\frac{v^2\tau}{2D}\left(\sqrt{4D^2k^2/v^2+1}+1\right)}}{k^2\sqrt{4D^2k^2 + v^2}} \tag{S30b}$$

$$B_3(k) = -\frac{a\rho}{D\sqrt{2\pi}} \frac{v + \sqrt{4D^2k^2 + v^2}}{k^2\sqrt{4D^2k^2 + v^2}} \tag{S30c}$$

$$B_4(k) = \frac{a\rho}{D\sqrt{2\pi}} \frac{\left(v - \sqrt{4D^2k^2 + v^2}\right)\left(1 - e^{-\frac{v^2\tau}{2D}\left(\sqrt{4D^2k^2/v^2+1}-1\right)}\right)}{k^2\sqrt{4D^2k^2 + v^2}} \tag{S30d}$$

Inverse Fourier transforming at $z = 0$ gives the real-space concentration and the following self-consistency relationship for the wave speed:

$$C_{\text{th}} = \frac{1}{\sqrt{2\pi}} \int_{-\infty}^{\infty} dk\ B_4(k), \tag{S31}$$

which simplifies to $C_{\text{th}} = 2a\rho/\pi v$ for $\tau \gg D/v^2$.

We can compare $\tau$ to the characteristic time $\tau_c = D(\pi C_{\text{th}}/2a\rho)^2$, which is equal to $D/v^2$ in the limit of continuous emission. Numerical solution of (S31) reveals that there is no self-consistent solution to (S31) until about $\tau \approx 1.53\tau_c$, at which point $v \approx 0.6 \times 2a\rho/\pi C_{\text{th}}$; $\tau \approx 1.53\tau_c$ is larger than the minimum initiation time of $t_{\text{min,3D}} = 4\tau_c/\pi$ (Fig. 2 C, Section 7). Thus, an initial pulse of length $4\tau_c/\pi < \tau < 1.53\tau_c$ from cells within an initial signaling radius $r_i$ can cause neighboring cells to exceed $C_{\text{th}}$, but cannot trigger a wave-like solution asymptotically.

#### Continuous emission plus decay with cells in 2D and diffusion in 3D

We can add signaling molecule decay to the embedded system dynamics by adding a term of $-\gamma C(\tilde{r}, k)$ to (S14) with $\gamma$ the signaling molecule decay rate. Going through the same exercise yields:

$$C(\tilde{r} < 0, k) = \sqrt{\frac{2}{\pi}} \frac{a\rho}{\gamma + k^2D} - \frac{a\rho}{\sqrt{2\pi}} \frac{\sqrt{4D^2k^2/v^2 + 4D\gamma/v^2 + 1} + 1}{(Dk^2 + \gamma)\sqrt{4D^2k^2/v^2 + 4D\gamma/v^2 + 1}} e^{\frac{\tilde{r}v}{2D}\left(-1+\sqrt{4D^2k^2/v^2+4D\gamma/v^2+1}\right)} \tag{S32a}$$

$$C(\tilde{r} > 0, k) = \frac{a\rho}{\sqrt{2\pi}} \frac{\sqrt{4D^2k^2/v^2 + 4D\gamma/v^2 + 1} - 1}{(Dk^2 + \gamma)\sqrt{4D^2k^2/v^2 + 4D\gamma/v^2 + 1}} e^{-\frac{\tilde{r}v}{2D}\left(1+\sqrt{4D^2k^2/v^2+4D\gamma/v^2+1}\right)} \tag{S32b}$$

with

$$C_{\text{th}} = \frac{a\rho}{\pi\sqrt{D\gamma}} \arcsin\left[\left(1 + v^2/4D\gamma\right)^{-1/2}\right] \tag{S33}$$

as the parameter relationship obtained after an inverse Fourier transform at $z = 0$ and $x = 0$. This gives a profile that propagates as a pulse (Fig. 1B).

In the limit $\gamma \ll v^2/D$, we recover the familiar expression $C_{\text{th}} = 2a\rho/\pi v$. Again, as when we accounted for signaling molecule decay with cells in 2D and diffusion in 2D, the wave speed approaches zero, but with $\gamma = (\pi/4)^2/\tau_c$ where $\tau_c = D(\pi C_{\text{th}}/2a\rho)^2$ (Fig. 2 D).

### 4 Finite extracellular medium

We consider now what happens when one does not lie in the extreme cases of an extracellular medium of thickness $h \gg D/v$ or $h \ll D/v$. To examine the case of arbitrary thickness $h$, we turn to the method of images (Fig. 3A).

Our boundary conditions for the extracellular medium require the concentration to obey $\partial c/\partial z = 0$ at $z = 0, h$ – signaling molecules cannot diffuse downward or upward once they reach the boundaries. By invoking the uniqueness theorem, we know that if we can find an arrangement of "image cells" – each emitting diffusible signaling molecule at rate $a$ – that satisfies these boundary conditions, then this arrangement of cells gives the *unique* solution for the concentration profile inside the extracellular medium. In our case, to satisfy the boundary condition above, we have image cells at $z = \pm 2jh$ for $j = 1, 2, \ldots$.

This means that we can find the concentration profiles simply by adding up the contributions from many discrete sources. Given this knowledge, we seek a relationship like (S11) or (S16) but for arbitrary $h$. To do so, we use Green's function integration and the fact that we can analyze the asymptotic dynamics of cells in 1D to deduce the asymptotic dynamics of cells in 2D, as previously shown.

The concentration – as measured at $(r, z = 0, t)$ – of a burst-like emission by a single point-like source at $(R, 2jh, T)$ is given by the Green's function:

$$G(r, z = 0, t; R, 2jh, T) = \frac{e^{-\frac{(r-R)^2 + (2jh)^2}{4D(t-T)}}}{2\pi D(t-T)}, \tag{S34}$$

so the Green's function of a single point-like source in a finite-thickness extracellular medium is (Fig. 3A):

$$G_h(r, z = 0, t; R, T) = \sum_{j=-\infty}^{\infty} G(r, z = 0, t; R, 2jh, T) = \frac{e^{-\frac{(r-R)^2}{4D(t-T)}}}{2\pi D(t-T)} \left(1 + 2\sum_{j=1}^{\infty} e^{\frac{-(jh)^2}{D(t-T)}}\right). \tag{S35}$$

We assume a traveling wave solution at speed $v$ meaning that the concentration $C_{\text{th}} = c(r = vt, z = 0, t \to \infty)$ is, for a density of cells $\rho$ emitting with rate $a$, given by:

$$C_{\text{th}} = c(vt, 0, t \to \infty) = a\rho \lim_{t\to\infty} \int_0^{vt} dR \int_{R/v}^{t} dT \; G_h(vt, 0, t; R, T) =$$
$$\frac{a\rho}{2\pi D} \int_{-\infty}^{0} d\tilde{R} \int_0^{-\tilde{R}/v} \frac{d\tilde{t}}{\tilde{t}} e^{-\frac{\tilde{R}^2}{4D\tilde{t}}} \left(1 + 2\sum_{j=1}^{\infty} e^{-\frac{(jh)^2}{D\tilde{t}}}\right) \tag{S36}$$

with the substitutions $\tilde{t} = t - T$ and $\tilde{R} = R - vt$. This yields:

$$C_{\text{th}} = \frac{2a\rho}{\pi v} \left(1 + 2\sum_{j=1}^{\infty} \int_0^{-\infty} dx \; \text{Ei}\left[\frac{1}{x}\left(\frac{jhv}{2D}\right)^2 + x\right]\right) = \frac{2a\rho}{\pi v} \sum_{j=-\infty}^{\infty} \int_0^{-\infty} dx \; \text{Ei}\left[\frac{1}{x}\left(\frac{jhv}{2D}\right)^2 + x\right] \tag{S37}$$

for $x = v\tilde{R}/4D$ and Ei[.] the exponential integral function. For $h \gg D/v$, (S37) reduces to (S16) because the term $\sum_{j=1}^{\infty} \cdots \approx 0$. Meanwhile, for $h \ll D/v$, the sum over $j$ can be turned into an integral, giving the familiar thin extracellular medium relationship, (S11).

We emphasize that (S37) provides a universal relationship between threshold concentration, wave speed, cell density, and signaling molecule emission rate for any extracellular medium thickness. By dividing both sides of (S37) by $a\rho h/\pi D$, we arrive at a relationship between a non-dimensionalized threshold concentration, $\pi C_{\text{th}} D/a\rho h$, and a non-dimensionalized wave speed, $vh/2D$:

$$\frac{\pi C_{\text{th}} D}{a\rho h} = \frac{2D}{vh} \sum_{j=-\infty}^{\infty} \int_0^{-\infty} dx \; \text{Ei}\left[\frac{1}{x}\left(\frac{jhv}{2D}\right)^2 + x\right]. \tag{S38}$$

We plot this relationship in Fig. 3B and see that (S38) is an interpolation between the thin ($h \ll D/v$) and thick ($h \gg D/v$) extracellular medium limits.

### 5 Asymptotic wave dynamics with Hill function activation

#### Numerical solutions show traveling waves

As shown in the main text, making the change from a Heaviside function source term to an order-$n$ Hill function ($\Theta[c - C_{\text{th}}] \to c^n/(c^n + C_{\text{th}}^n)$) in (S1) preserves the scaling relationships (S11) and (S16) with a constant factor as long as the new source terms give traveling wave solutions. We have found numerically that $n \geq 1$ Hill functions indeed give traveling wave solutions, with $n = 1, 2, 3$ shown for thin and thick extracellular media in Fig. 4.

To find these solutions, we numerically solved (S1) with a Hill function source term for cells in 1D with diffusion in one or two dimensions. We used $D = 10^{-10}$ m$^2$/s and $v_\Theta = 2$ $\mu$m/s with the threshold concentration determined by (S16) and (S11) with $v \to v_\Theta$. In this way, we could compare $v_n$ – the wave speed given by the order-$n$ Hill function – to the Heaviside wave speed $v_\Theta$. For our numerics, we imposed a maximum size step of $D/10v_\Theta$ for the spatial dimension where the cells live, a maximum step size of $D/30v_\Theta$ for the added spatial dimension (when modeling diffusion in 2D), and a maximum time step of $D/10v_\Theta^2$. These step sizes give convergence of the information wave fronts, which we define as the curves $r_c(t)$ such that $c(r_c(t), t) = C_{\text{th}}$. We simulated times $t_{\max} \leq 25D/v_\Theta^2$ using Mathematica's "NDSolveValue" function and, when modeling diffusion in 2D, replaced the delta function in (S1) with a Gaussian of width $D/10v_\Theta$. This substitution gives a wave speed (according to an inverse Fourier transform of (S15b) at $\tilde{r}, z = 0$) of $v/v_\Theta \approx 0.95$. The initiating colony is of size $r_i = 2D/v_\Theta$, and we assume all cells in the initiating colony signal at the maximal rate $a$.

To find the wave speeds noted in (4), we found the location of the wave front, $r_c(t)$, and fit a line to the region of $24D/v_\Theta^2 \leq t \leq 25D/v_\Theta^2$. As shown in 4, for $n \geq 2$, the wave speeds are very close to $v_\Theta$ with significant deviation only for $n = 1$. Even the concentration profiles are in good agreement with the Heaviside solution for $n \geq 2$.

#### Connection to Fisher Waves

When the dimensionality of the cell distribution matches the diffusive dimensionality and cell activation is described by the $n = 1$ Hill function, one can find the wave speed by using a modified version of the analysis pioneered by Fisher and Kolmogorov *et al.* [6, 7]. To see this, we first consider a modified version of (S3c) with order-$n$ Hill function activation:

$$0 = D\frac{\partial^2 c}{\partial \tilde{r}^2} + v\frac{\partial c}{\partial \tilde{r}} + a\rho\frac{c^n}{c^n + C_{\text{th}}^n}. \tag{S39}$$

The new activation term is mathematically obnoxious as it no longer has a simple spatial interpretation; with a Heaviside function, for example, one can turn a term like $\Theta[c - C_{\text{th}}]$ into a simple function of $\tilde{r}$: $\Theta[c - C_{\text{th}}] = \Theta[-\tilde{r}]$. This has an important consequence: instead of solving two differential equations with constant source terms and matching boundary conditions (as we did for the Heaviside emission), we must now solve a single differential equation with a difficult non-linear source term. We note that (S39) is, in the limit $n \to \infty$, equivalent to a relay with Heaviside activation.

However, there is a distinct advantage to the new source term: it is one-to-one in $c$. Thus, we may make the substitution $f = \frac{c^n}{c^n + C_{\text{th}}^n} \implies c = C_{\text{th}}\left(\frac{f}{1-f}\right)^{1/n}$, then plug this into eq. (S39) to yield (after some rearrangement):

$$0 = v\frac{\partial f}{\partial \tilde{r}} + D\left[\frac{\partial^2 f}{\partial \tilde{r}^2} + \frac{1 + (2f - 1)n}{nf(1 - f)}\left(\frac{\partial f}{\partial \tilde{r}}\right)^2\right] + \frac{na\rho}{C_{\text{th}}}f^{2-1/n}(1 - f)^{1+1/n}. \tag{S40}$$

(S40) looks a lot like the traditional Fisher equation,

$$0 = v\frac{\partial f}{\partial \tilde{r}} + D\frac{\partial^2 f}{\partial \tilde{r}^2} + \frac{na\rho}{C_{\text{th}}}f(1 - f) \tag{S41}$$

in that it has a source term that goes to zero at $f \to \{0, 1\}$, a term $D\frac{\partial^2 f}{\partial^2 \tilde{r}}$, and a term $v\frac{\partial f}{\partial \tilde{r}}$. We therefore take Fisher's approach [6] and think of gradients of $f$ as functions of $f$ rather than $x$. As such, we define

$F(f) = \frac{\partial f}{\partial x}$, which allows us to make the substitution $\frac{\partial^2 f}{\partial x^2} = \frac{\partial F}{\partial x} = \frac{\partial f}{\partial x}\frac{\partial F}{\partial f} = F\frac{\partial F}{\partial f}$. This is valid under the assumption that the concentration profiles are monotone decreasing, which is seen to be the case in Fig. 4. Under all of the above, we get:

$$0 = vF + D\left[F\frac{\partial F}{\partial f} + \frac{1+(2f-1)n}{nf(1-f)}F^2\right] + \frac{na\rho}{C_{\text{th}}}f^{2-1/n}(1-f)^{1+1/n} \tag{S42}$$

which gives us a non-linear ODE for $F$. Of particular note are the boundary conditions for $F$. Namely, $F(0) = F(1) = 0$, which is to say that cells well inside of the wave front are all emitting at their maximal rate since $c \gg C_{\text{th}}$ and that cells well beyond the wave front are not emitting at all. Thus, there is no spatial dependence on the cellular activation, $f = c^n/\left(c^n + C_{\text{th}}^n\right)$, in these regions.

Next, we turn to the traditional method of examining the $f \to 0$ limit. As $F \to 0$, (S42) becomes, to lowest order in $f$ and assuming $F \approx -\lambda f^\beta$,

$$0 = -\lambda v + D\left[\lambda^2\beta f^{\beta-1} + \frac{1-n}{n}\lambda^2 f^{\beta-1}\right] + \frac{na\rho}{C_{\text{th}}}f^{2-1/n-\beta}. \tag{S43}$$

where $\lambda > 0$. One can only obtain a self-consistency relationship between $v$ and $C_{\text{th}}/a\rho$ if $\beta = 2 - 1/n$. Otherwise, $f^{2-1/n-\beta}$ diverges or goes to zero. With this choice, (S43) becomes:

$$n = 1\colon\ 0 = -\lambda v + D\lambda^2/n + \frac{a\rho}{C_{\text{th}}} \implies \lambda = \frac{1}{2D}\left(v \pm \sqrt{v^2 - \frac{4a\rho D}{C_{\text{th}}}}\right) \tag{S44a}$$

$$n > 1\colon\ 0 = -\lambda v + \frac{na\rho_0}{C_{\text{th}}} + \mathcal{O}(f^{1-1/n}) \implies \lambda = \frac{na\rho}{vC_{\text{th}}} \tag{S44b}$$

where in (S44b) we have taken the $f \to 0$ limit. To get a wave speed, $v$, out of (S44a), we demand that the quantity under the square root be non-negative, which ensures that $\lambda > 0$ is a real number as assumed. This means $v \geq 2\sqrt{a\rho D/C_{\text{th}}} = 2v_\Theta$ – a bound that is very similar to conventional Fisher waves. In the case of Fisher waves, the minimum wave speed is selected for [6, 7]; the same is true here, as the minimum wave speed $v = 2\sqrt{a\rho D/C_{\text{th}}}$ is what one finds after numerically solving the 1D dynamics (Fig. 4A). In contrast, for $n > 1$, this method yields no such wave speed bound.

### 6 Assessing the validity of a continuum analysis

Next, we consider the validity of a continuum analysis like (S3b) for studying asymptotic wave dynamics. To do so, we compare our continuum wave speed relationships, (S11) and (S16), to a simple model of discrete cells on a lattice in 1D (Fig. 5A). We refer to this as the discrete lattice model. We briefly discuss the results of this lattice model below, and show that it agrees with the continuous model when the separation between cells, $d$, is much less than the characteristic length $D/v$. This heuristic also holds for cells in two and three dimensions, and for cells scattered randomly according to a Poisson process.

To start, we first calculate the three-dimensional concentration (SI units of $1/\text{m}^3$) generated by a continuously emitting point source emitting at a rate $a$ at a distance $x$ after a time $t$. For diffusion in $m$ dimensions, we will refer to this concentration as $c_{\bullet,m}(x,t)$. For diffusion in $m = 2$ dimensions, we will consider a semi-infinite environment in order to recapitulate (S16), which holds for cells in one (two) dimensions with diffusion in a semi-infinite two-dimensional (three-dimensional) space. These relationships are:

$$c_{\bullet,1}(x,t) = \frac{a}{h^2}\int_0^t dT\, \frac{e^{-x^2/4DT}}{(4\pi DT)^{1/2}} = \frac{a}{h^2}\sqrt{\frac{t}{\pi D}}\left(e^{-x^2/4Dt} - \sqrt{\frac{\pi x^2}{4Dt}}\text{erfc}\sqrt{\frac{x^2}{4Dt}}\right) \tag{S45a}$$

$$c_{\bullet,2}(x,t) = \frac{2a}{h}\int_0^t dT\, \frac{e^{-x^2/4DT}}{4\pi DT} = -\frac{a}{2\pi hD}\text{Ei}\left(-\frac{x^2}{4Dt}\right) \tag{S45b}$$

with erfc[.] the complementary error function and Ei[.] the exponential integral. We have assumed one-dimensional diffusion takes place in a narrow channel with cross-sectional area $h^2$ and two-dimensional diffusion takes place in an extracellular medium of thickness $h$.

Next, we assume the cells in this lattice model perform a signaling relay with Heaviside activation: once the local concentration exceeds a threshold $C_{\mathrm{th}}$, they participate in the signaling molecule emission at a rate $a$. If the resulting wave speed is $v$, then a cell at a distance $\tilde{x} = -jd$ from the wave front has been emitting for a time $jd/v$. That cell then creates a concentration $c_{j,m} = c_{\bullet,m}(jd, jd/v)$ at $(\tilde{x}, t) = (0, 0)$. The full concentration at the wave front is $C_{\mathrm{th}}$ by definition, and it is equal to the sum of the concentrations created by all cells behind the wave front. This gives us the following self-consistency relationships between the threshold concentration, $C_{\mathrm{th}}$; wave speed, $v$; diffusion constant, $D$; and cell separation $d$:

$$\text{diff. in 1D: } C_{\mathrm{th}} = \sum_{j=1}^{\infty} c_{j,1} = \frac{a}{vh^2} \sum_{j=1}^{\infty} \sqrt{j\frac{vd}{\pi D}} \left[ e^{-jvd/4D} - \sqrt{j\frac{\pi vd}{4D}} \mathrm{erfc}\left( \sqrt{j\frac{vd}{4D}} \right) \right] \tag{S46a}$$

$$\text{diff. in 2D: } C_{\mathrm{th}} = \sum_{j=1}^{\infty} c_{j,2} = -\frac{a}{2\pi h D} \sum_{j=1}^{\infty} \mathrm{Ei}\left( -j\frac{vd}{4D} \right). \tag{S46b}$$

(S46a) and (S46b) provide relationships analogous to (S11) and (S16). In fact, in the limit $vd/4D \ll 1$, the sums in these relationships are well-approximated by an integral over $j$ from $j = 0$ to $\infty$. In this limit, with $\rho = 1/d$, (S46a) becomes $C_{\mathrm{th}} = a\rho D/h^2 v^2$, the one-dimensional analog of (S11); similarly, (S46b) simplifies to $C_{\mathrm{th}} = 2a\rho/\pi h v$, the one-dimensional analog of (S16). (One can turn (S46b) into an integral from $j = 0$ to $\infty$. The integrand diverges at $j = 0$, but is still integrable because the divergence is logarithmic.) Therefore, we can see that the continuum limit is valid when the separation between cells is $d \ll 4D/v$. We have shown the approach to the continuous theory limit in Fig. 5B/C.

### 7 Initiation dynamics

In this section, we demonstrate the initiation time relationships discussed in the main text using Green's function integration. To do so, we write down the Green's functions $G_{n,m}(r, t; R, T)$ describing diffusion of molecules in $m$ dimensions released by cells in $n$ dimensions at $(R, T)$ and measured by cells at $(r, t)$. For $n \neq m$, we assume a semi-infinite environment. For cells in 1D and diffusion in 1D, we assume a narrow channel of width $h$ in both dimensions perpendicular to the channel. For cells in 1D or 2D and diffusion in 2D, we assume an extracellular medium of thickness $h$. We calculate the Green's functions for cells in two dimensions by integrating over a ring of diffusive sources at radius $R$; we calculate the Green's functions for cells in three dimensions by integrating over a shell of diffusive sources at radius $R$. Below, $I_0[.]$ is the zeroth $I$-Bessel function and $\sinh[.]$ is the hyperbolic sine function. The Green's functions are

$$G_{1,1}(r, t; R, T) = e^{-(r-R)^2/4D(t-T)}/h^2\sqrt{4\pi D(t-T)} \tag{S47a}$$

$$G_{1,2}(r, t; R, T) = e^{-(r-R)^2/4D(t-T)}/2\pi h D(t-T) \tag{S47b}$$

$$G_{2,2}(r, t; R, T) = R\; I_0\left[rR/2D(t-T)\right] e^{-(r^2+R^2)/4D(t-T)}/2hD(t-T) \tag{S47c}$$

$$G_{2,3}(r, t; R, T) = R\; I_0\left[rR/2D(t-T)\right] e^{-(r^2+R^2)/4D(t-T)}/2\sqrt{\pi}D^{3/2}(t-T)^{3/2} \tag{S47d}$$

$$G_{3,3}(r, t; R, T) = R \sinh\left[rR/2D(t-T)\right] e^{-(r^2+R^2)/4D(t-T)}/r\left[\pi D(t-T)\right]^{1/2}. \tag{S47e}$$

One-by-one, we study the initiation time for these systems by studying the self-consistency relationship for the threshold concentration $C_{\mathrm{th}}$ and initiation time $t_{\mathrm{init}}$ for a given initial signaling colony of size $r_i$:

$$C_{\mathrm{th}} = a\rho \int_0^{t_{\mathrm{init}}} dT \int_0^{r_i} dR\; G_{n,m}(r_i, t_{\mathrm{init}}; R, T) = a\rho \int_0^{t_{\mathrm{init}}} dT \int_0^{r_i} dR\; G_{n,m}(r_i, 0; R, -T) \tag{S48}$$

where the logic here is that the signaling wave initiates when the threshold concentration at the edge of the initial signaling colony exceeds $C_{\mathrm{th}}$. At $t_{\mathrm{init}}$, cells outside the colony participate in the signaling and birth a diffusive wave with dynamics we have already studied extensively. This scenario assumes the cells do not move – that the cell density is fixed. In the case of neutrophils, this means that we are ignoring the possibility that a cell initially located off the target randomly encounters the target and starts signaling. Unlike the asymptotic dynamics, the difference in diffusive and cell dimension is not the salient parameter for understanding diffusive wave initiation. Rather, the initiation dynamics are determined solely by the dimension of the diffusive environment.

We now seek to derive the equations in the main text which show the relationship between the wave initiation time $t_{\text{init}}$ and initial signaling colony size $r_i$ for various system dimensionalities when $r_i \gg D/v$ or $r_i \ll D/v$. The full relationships of $t_{\text{init}}$ versus $r_i$ for various system dimensionalities are plotted in Fig. 3 of the main text.

#### Initiation with cells in 1D and diffusion in 1D

With cells and diffusion in 1D, we can perform the integral (S48) directly (for cells in 1D, we consider the bounds on the integral over $R$ to be $-r_i$ to $r_i$) and get a closed form relationship:

$$C_{\text{th}} = \frac{a\rho r_i^2}{h^2 D}\left[\left(Dt_i/\pi r_i^2\right)^{1/2} e^{-r_i^2/Dt_i} - 1 + \left(1 + Dt_i/2r_i^2\right)\text{erf}\left(r_i^2/Dt_i\right)^{1/2}\right]. \tag{S49}$$

In the limit where $r_i \gg Dt_i$, we get that $t_i = 2D/v^2$ by using the asymptotic relationship (S11). This is the minimum initiation time, $t_{\text{min},1,1}$

$$t_{\text{min},1,1} = 2D/v^2 \tag{S50}$$

and it tells us that $r_i \gg Dt_i$ is equivalent to $r_i \gg D/v$ – we can appeal to the natural length and time scales from our asymptotic analysis. We will soon see that this is also the minimum initiation time for cells in 2D with diffusion in 2D and cells in 3D with diffusion in 3D; this is the case because, as in the asymptotic analysis, we've essentially ignored the curvature of the target when calculating the $r_i \gg D/v$ initiation time.

In the opposite limit – $r_i \ll D/v$ – we can Taylor Expand (S49) and get

$$r_i \ll D/v : t_{\text{init}} \approx (\pi D/4v^2)(D/vr_i)^2, \tag{S51}$$

thus validating our equations in the main text.

#### Initiation with cells in 1D and diffusion in 2D

Next, we consider the self-consistency equation (S48) with $n = 1, m = 2$. As before, we first consider the limit of $r_i \gg Dv$ and recover (through (S16)):

$$t_{\text{min},1,2} = 4D/\pi v^2 \tag{S52}$$

while for $r_i \ll Dv$,

$$r_i \ll D/v : \log\left(Dt_{\text{init}}/r_i^2\right) \approx 2D/vr_i. \tag{S53}$$

#### Initiation with cells in 2D and diffusion in 2D

Moving on, we consider the case in the main text of cells in 2D with diffusion in 2D. To perform the integration of (S48) in this case, it is easiest to rewrite the Bessel function in (S47c) in integral form, then integrate first over time. With $r_i \gg D/v$, such an analysis gives a minimum initiation time of

$$t_{\text{min},2,2} = 2D/v^2. \tag{S54}$$

In the opposite limit of $r_i \ll D/v$,

$$r_i \ll D/v : \log(4Dt_{\text{init}}/r_i^2) \approx (2D/vr_i)^2 \tag{S55}$$

as noted in the main text.

##### Initiation with cells in 2D and diffusion in 3D

Now, we will see that diffusive waves do not always initiate in 3D environments. We consider the integral in (S48), but take $t_{\text{init}} \to \infty$ which gives us a maximum concentration $C_{\text{max},2,3}$ at $r = r_i$ of:

$$C_{\text{max},2,3} = 2a\rho r_i/\pi D \tag{S56}$$

Thus, by (S16), if $r_i < D/v$, $C_{\text{max},2,3} < C_{\text{th}}$ and the signaling wave cannot initiate. Examining (S48) for $r_i \approx D/v$ reveals

$$r_i \approx D/v : t_{\text{init}} \approx \left(\pi D/16v^2\right)(vr_i/D)^4(vr_i/D - 1)^{-2} \tag{S57}$$

while

$$t_{\text{min},2,3} = 4D/\pi v^2. \tag{S58}$$

##### Initiation with cells in 3D and diffusion in 3D

Finally, we consider initiation with cells in 3D. Again, we consider the limit $t_{\text{init}} \to \infty$ in (S48) to get:

$$C_{\text{max},3,3} = a\rho r_i^2/3D. \tag{S59}$$

Thus, waves do not initiate below a critical $r_i$. However, here, the critical value is $r_i = \sqrt{3}D/v$. As with 1D cells/diffusion and 2D cells/diffusion, we recover

$$t_{\text{min},3,3} = 2D/v^2 \tag{S60}$$

in the limit $r_i \gg D/v$.

### 8 Wave initiation with Hill function activation

We now characterize the effect of Hill function-like activation on the signaling wave initiation, as we have done already for asymptotic dynamics. To do so, we consider a simple situation: cells within a volume of size $r_i$ signal with some rate $a$ while neighboring cells outside of the initial signaling volume (i.e., with $r > r_i$) signal with a concentration-dependent rate of $ac^n/(c^n + C_{\text{th}}^n)$. These calculations give us an idea of how sensitive the initiation dynamics are to the details of cell activation. As we will show, the initiation dynamics with Hill function activation are a good approximation of the initiation dynamics for Heaviside activation when for relatively small $n$. One can imagine that such a situation may be relevant in, e.g., the neutrophil swarming experiments presented in the main text [8]. Here, cells in direct contact with a foreign protein begin signaling their neighbors, which respond to the presence of the signaling molecule by participating in the emission themselves. This analysis again treats the cell distribution as static and ignores the possibility that neutrophils may randomly encounter the target.

##### Wave initiation with Hill function activation, cells in 1D, and diffusion in 1D

For cells in 1D with diffusion in 1D, the scenario described above can be described with the following equation of motion:

$$\frac{\partial c}{\partial t} = D\frac{\partial^2 c}{\partial r^2} + \frac{a\rho}{h^2}\Theta[r_i - |r|] + \frac{a\rho}{h^2}\Theta[|r| - r_i]\frac{c^n}{c^n + C_{\text{th}}^n}. \tag{S61}$$

One can non-dimensionalize (S61) by dividing all the concentration scales by $C_{\text{th}}$, dividing all the length scales by $l_c = \sqrt{h^2 C_{\text{th}} D/a\rho}$, and dividing all the time scales by $\tau_c = h^2 C_{\text{th}}/a\rho$, thusly arriving at

$$\frac{\partial(c/C_{\text{th}})}{\partial(t/\tau_c)} = \frac{\partial^2(c/C_{\text{th}})}{\partial(r/l_c)^2} + \Theta\left[\frac{r_i - |r|}{l_c}\right] + \Theta\left[\frac{|r| - r_i}{l_c}\right]\frac{(c/C_{\text{th}})^n}{(c/C_{\text{th}})^n + 1}, \tag{S62}$$

which shows that $l_c = \sqrt{h^2 C_{\rm th} D/a\rho}$ is the relevant length scale and $\tau_c = h^2 C_{\rm th}/a\rho$ is the relevant time scale. (In the $n \to \infty$ limit of Heaviside activation, $\tau_c = D/v^2$ and $l_c = D/v$.) As with Heaviside activation, we refer to the initiation time $t_{\rm init}$ as the time at which $c(r_i, t_{\rm init}) = C_{\rm th}$. To find $t_{\rm init}$, we numerically solve (S61) using the methods discussed in Section 5. This gives the relationship of $t_{\rm init}/\tau_c$ as a function of $r_i/l_c$ and $n$ shown in Fig. 6A. As seen in Fig. 6A, even low-order ($n = 1, 2, 3, 5, 10$) Hill functions exhibit relatively large (compared to $\tau_c$) initiation times for $r_i \ll l_c$.

#### Wave initiation with Hill function activation, cells in 2D, and diffusion in 2D

Next, we study the initiation dynamics above with cells and diffusion in two dimensions. Here, the dynamics are governed by the following equation of motion:

$$\frac{\partial c}{\partial t} = D\left(\frac{\partial^2 c}{\partial r^2} + \frac{1}{r}\frac{\partial c}{\partial r}\right) + \frac{a\rho}{h}\Theta[r_i - r] + \frac{a\rho}{h}\Theta[r - r_i]\frac{c^n}{c^n + C_{\rm th}^n}. \tag{S63}$$

Now that we are considering the initiation dynamics, we must include terms like $D(\partial c/\partial r)/r$, which we could previously neglect in our asymptotic analysis of cells in 2D with diffusion in 2D. The curvature of the initial signaling colony matters when calculating initiation times. Note that (S63) can be non-dimensionalized in the same spirit as (S62) with characteristic length scale $l_c = \sqrt{hC_{\rm th}D/a\rho}$ and characteristic time scale $\tau_c = hC_{\rm th}/a\rho$. As with cells and diffusion in 1D, we numerically solve (S63) using the methods discussed in Section 5 to find $t_{\rm init}$ such that $c(r_i, t_{\rm init}) = C_{\rm th}$. Here again, we see that even low-order ($n = 2, 3, 5, 10$) Hill functions exhibit relatively large (compared to $\tau_c$) initiation times for $r_i \ll l_c$.

#### Wave initiation with Hill function activation, cells in 3D, and diffusion in 3D

Finally, we study signaling wave initiation properties of a 3D environment by studying wave initiation with cells and diffusion in 3D. To do so, we numerically solve the 3D analog of (S63),

$$\frac{\partial c}{\partial t} = D\left(\frac{\partial^2 c}{\partial r^2} + \frac{2}{r}\frac{\partial c}{\partial r}\right) + a\rho\Theta[r_i - r] + a\rho\Theta[r - r_i]\frac{c^n}{c^n + C_{\rm th}^n}, \tag{S64}$$

which can be non-dimensionalized in the same spirit as (S62), but with characteristic length scale $l_c = \sqrt{C_{\rm th}D/a\rho}$ and characteristic time scale $\tau_c = C_{\rm th}/a\rho$. We solve (S64) using the methods discussed in Section 5 to find $t_{\rm init}$ such that $c(r_i, t_{\rm init}) = C_{\rm th}$. This gives the numerically determined relationship of $t_{\rm init}$ as a function of $r_i$ and $n$ shown in Fig. 6C. We see that even low-order ($n = 2, 3$) Hill functions exhibit relatively large (compared to $\tau_c$) initiation times for $r_i \ll l_c$. Larger yet Hill functions ($n = 5, 10$) can lead to very large initiation times (compared to $\tau_c$) even for $r_i \approx l_c$.

In fact, $n > 3$ activation functions can result in initiation failures when $a\rho r_i^2/3DC_{\rm th} \ll 1$. To see that this is the case, we treat (S64) in the steady state ($\partial c/\partial t = 0$) and use a perturbative analysis, assuming $c^n/(c^n + C_{\rm th}^n) \ll 1$, in which case $c^n/(c^n + C_{\rm th}^n) \approx (c/C_{\rm th})^n$. In such a situation, we can write $c(r) \approx c_0(r) + c_1(r)$ as the sum of a dominant contribution $c_0$ that is generated by cells within $r_i$ and satisfies

$$0 = D\left(\frac{\partial^2 c_0}{\partial r^2} + \frac{2}{r}\frac{\partial c_0}{\partial r}\right) + a\rho\Theta[r_i - r] \tag{S65}$$

and a small correction $c_1$ that is generated by cells beyond $r_i$ and obeys

$$0 = D\left(\frac{\partial^2 c_1}{\partial r^2} + \frac{2}{r}\frac{\partial c_1}{\partial r}\right) + a\rho\frac{c_0^n}{c_0^n + C_{\rm th}^n}\Theta[r - r_i] \approx D\left(\frac{\partial^2 c_1}{\partial r^2} + \frac{2}{r}\frac{\partial c_1}{\partial r}\right) + a\rho\left(\frac{c_0}{C_{\rm th}}\right)^n\Theta[r - r_i]. \tag{S66}$$

We can solve (S65) directly to arrive at:

$$c_0(r < r_i) = \frac{a\rho}{2D}\left(r_i^2 - r^2/3\right),\ c_0(r > r_i) = \frac{a\rho r_i^3}{3Dr} \tag{S67}$$

where the form of $c_0(r > r_i)$ is reminiscent of solving for the potential of a uniformly charged sphere in electrostatics [9]. If $a\rho r_i^2/3DC_{\rm th} = \epsilon \ll 1$, we can calculate the perturbation $c_1$. By (S66), $c_1$ obeys

$$0 = D\left(\frac{\partial^2 c_1}{\partial r^2} + \frac{2}{r}\frac{\partial c_1}{\partial r}\right) + a\rho\left(\frac{c_0}{C_{\rm th}}\right)^n \Theta[r - r_i] = D\left(\frac{\partial^2 c_1}{\partial r^2} + \frac{2}{r}\frac{\partial c_1}{\partial r}\right) + a\rho\left(\frac{\epsilon r_i}{r}\right)^n \Theta[r - r_i] \quad \text{(S68)}$$

so that

$$c_1(r < r_i) = \frac{a\rho r_i^2 \epsilon^n}{D(n-2)},\ c_1(r > r_i) = \frac{a\rho\epsilon^n}{D(n-3)}\left[\frac{r_i^3}{Dr} + \frac{r^2}{n-2}\left(\frac{r_i}{r}\right)^n\right]. \quad \text{(S69)}$$

For $c_1$ to be a sensible perturbative correction, we require it to be positive (since we are adding source terms to $c_0$ to get $c_1$) and much smaller than $c_0$. This is the case when $n > 3$. In this limit, it is smaller than $c_0$ by roughly a factor of $\epsilon^n$ – a very small correction. Thus, $n > 3$ activation functions can give steady-state concentration profiles that do not trigger waves in a three-dimensional diffusive environment.

In Fig. 6C, we can indeed see that small $n$ activation functions show less appreciable increases in the initiation time as $r_i$ decreases.

### 9 Green's function integration for information wave front finding

Having enumerated the Green's functions in the previous section, we now explain our numerical integration method used in the main text. To find the information wave front for cells in $n$ dimensions and diffusion in $m$ dimensions with continuous emission and Heaviside activation, one is looking for a curve $r_c(t)$ that defines the wave front: $C_{\rm th} = c(r_c(t), t)$. Thus, with an initial signaling colony of size $r_i$, one must solve the problem:

$$C_{\rm th} = a\rho \int_0^t dT \int_0^{\max[r_i, r_c(T)]} dR\ G_{n,m}(r_c(t), t; R, T). \quad \text{(S70)}$$

We do so by first finding the initiation time according to (S48), then finding $r_c(t)$ at discrete times, incrementing in steps of $\Delta t$ (we use $\Delta t = D/10v^2$ in the main text and Fig. 7, which gives convergence of the information wave front). We use linear interpolation between these points to define a continuous curve $r_c(t)$.

As an example of this method, we have plotted various information wave fronts in Fig. 7. These wave fronts assume different values of the diffusion constant $D$ and the threshold concentration $C_{\rm th}$ is fit to give the experimentally observed wave initiation time in ref. [8]. As we can see from the plot, there is a small range of values for $D$ for which one can construct an information wave front that agrees with the data (Fig. 7 A). These values of $D$ are consistent with the diffusion constant of small molecules like LTB4. Values of $D$ differing significantly from this range give information wave fronts that differ significantly from the experimentally observed wave front.

### 10 Simple diffusion model

In the main text, we showed the qualitative differences between a signaling relay, in which cells emit one type of signaling molecule in response to the local concentration of the same molecule, and a simple diffusive signaling model, in which cells within some volume signal surrounding cells, which do not participate in the signaling at all. Here, we explicitly calculate some of the properties of a simple diffusion model. We consider cells in 2D, with a region of cells of size $r_i$ at $z = 0$ in which the cells emit diffusible signaling molecules at rate $a$. In equation form,

$$\frac{\partial c}{\partial t} = D\left(\frac{\partial^2 c}{\partial r^2} + \frac{1}{r}\frac{\partial c}{\partial r} + \frac{\partial^2 c}{\partial z^2}\right) + a\rho\delta(z)\Theta[r_i - r] \quad \text{(S71)}$$

describes the concentration in space and time. To calculate concentrations, one can either propagate this equation directly or, as we do in the main text, integrate Green's functions in a manner similar to that described above. For cells in 2D with diffusion in 3D (assuming a semi-infinite environment), this gives

$$c(r,z=0,t) = a\rho \int_0^t dT \int_0^{r_i} dR\ G_{2,3}(r,0;R,-T) \tag{S72}$$

as the concentration at $z = 0$. One can take gradients according to $\partial c/\partial r$.

In the limit $r \gg r_i$, the signaling colony looks like a point source, meaning that (S72) can be simplified according to $\int dR\ G_{2,3} \approx r_i^2 e^{-r^2/4DT} / \left(4\sqrt{\pi} D^{3/2} T^{3/2}\right)$. Thus, we get that

$$r \gg r_i: \ c(r,z=0,t) \approx \frac{a\rho r_i^2}{2\sqrt{\pi} D^{3/2}} \int_0^t dT\ \frac{e^{-r^2/4DT}}{T^{3/2}} = \frac{a\rho r_i^2}{2Dr} \text{erfc} \left(\frac{r^2}{4Dt}\right)^{1/2} \tag{S73}$$

which, in the limit of $r^2 \gg Dt$, gives

$$r \gg r_i, \sqrt{Dt}: \ c(r,z=0,t) \approx \frac{a\rho r_i^2}{Dr} \text{erfc} \left(\frac{r^2}{4Dt}\right)^{1/2} \approx \frac{a\rho r_i^2}{r^2} \sqrt{\frac{t}{\pi D}}\ e^{-r^2/4Dt} \tag{S74}$$

which shows that the concentration profiles of simple diffusive models indeed have very shallow, Gaussian (with $1/r^2$ adjustments) tails.

Similarly, for cells in 2D with diffusion in 2D,

$$c(r,t) = \frac{a\rho}{h} \int_0^t dT \int_0^{r_i} dR\ G_{2,2}(r,0;R,-T) \tag{S75}$$

describes the concentrations. In the same limits ($r \gg r_i, \sqrt{Dt}$), we get

$$r \gg r_i, \sqrt{Dt}: \ c(r,t) \approx \frac{a\rho r_i^2 t}{hr^2} e^{-r^2/4Dt} \tag{S76}$$

as the concentration. Here again, we see that the concentration profiles are shallow Gaussian with $1/r^2$ adjustments. We plot the resulting gradients in this thin extracellular medium limit against the gradients from a comparable relay model in Fig. 8B.

### 11 Quantifying the effects of chemotaxis

To understand the effects that chemotaxis has on our model, we consider cells in 2D. The results below can be adopted to study cells in 3D or 1D, though the dimensionality of the cells has no effect on the asymptotic signaling wave speed.

We consider the same signaling motif as in the main text – that cells emit a diffusible molecule with rate $a$ once the local concentration of the same molecule exceeds $C_{\text{th}}$ – but now consider a time-varying density. Our model is a coarse-grained one; we study the case of cells moving toward the origin (radially inward) with a mean speed $u$ once the local concentration exceeds $C_{\text{th}}$. This is a toy model of neutrophil chemotaxis and is, of course, an approximation because the mean radial speed will depend on – among many other factors – the strengths of the gradients the cells use for chemotaxis. In full, for cells in any number of dimensions and within this model,

$$\frac{\partial c(\mathbf{r},t)}{\partial t} = D\nabla^2 c + a\rho(\mathbf{r},t)\Theta[c - C_{\text{th}}] \tag{S77a}$$

$$\frac{\partial \rho(\mathbf{r},t)}{\partial t} = \nabla \cdot (u\hat{r}\rho)\, \Theta[c(\mathbf{r},t) - C_{\text{th}}] \tag{S77b}$$

where $\hat{r}$ is the unit vector pointing radially outward. For cells in 2D,

$$\frac{\partial c(r,z,t)}{\partial t} = D\left(\frac{\partial^2 c}{\partial r^2} + \frac{1}{r}\frac{\partial c}{\partial r} + \frac{\partial^2 c}{\partial z^2}\right) + a\rho(r,t)\delta(z)\left(\Theta[c - C_{\text{th}}]\Theta[r - r_i] + \Theta[r_i - r]\right) \tag{S78a}$$

$$\frac{\partial \rho(r,t)}{\partial t} = \frac{u}{r}\frac{\partial\,(r\rho)}{\partial r}\Theta[c - C_{\text{th}}]\Theta[r - r_i] \tag{S78b}$$

which are the coupled equations we will study going forward. Note that we have included a source term for cells within the initial signaling colony of radius $r_i$.

We again assume a signaling wave propagates at outward with speed $v$. In this case, the cell density beyond the target is described by:

$$\rho(r_i < r < vt, t) = \rho_0 \frac{1 + ut/r}{(1 + u/v)^2} \tag{S79}$$

which one can derive by assuming an outward propagating wave with speed $v$, a group of inward chemotaxing cells with speed $u$, and an initially uniform density of cells $\rho_0$. To do so, we consider the signaling wave passing a cell at radius $R$ and time $t - T$. At a later time $t$, the cell initially at $R$ has moved inward a distance $uT$. Thus, the density at $r = R - uT$ and $t$ is $\rho(r,t) \sim \rho_0 R/r = \rho_0(1 + ut/r)/(1 + u/v)$. Integrating this density and demanding conservation of cell number gives (S79).

#### Effect on asymptotic wave speed relationships

Before numerically solving (S78a) and (S78b) to show the precise effects chemotaxis has on the concentration profiles, concentration profiles, and information wave fronts, we calculate the effect it has on the asymptotic wave speed.

To do so, we first show that only cells within $\approx D/v$ of the wave front contribute to the concentration at the wave front. To contribute to the concentration at the wave front, you need to be within about a diffusion length of it. If the wave front passed a time $t$ ago, that means being within $\delta r \approx \sqrt{Dt}$. However, we know that $t = \delta r/v$, so $\delta r \approx D/v$ – the characteristic length scale of diffusive waves.

As only cells within $\approx D/v$ of the wave front contribute to concentration at the wave front, we are considering cell densities on the order of:

$$\rho(r = vt - D/v, t) = \rho_0 \frac{1 + ut/r}{(1 + u/v)^2} \approx \rho_0 \frac{1 + ut/(vt - D/v)}{(1 + u/v)^2}. \tag{S80}$$

In the asymptotic regime of $vt \gg D/v$, we get

$$\rho \approx \rho_0/(1 + u/v) \tag{S81}$$

meaning the density of cells that contribute to the wave front propagation is approximately constant. Therefore, we may modify the analysis that lead to (S11) and (S16) to get two new asymptotic equations for the wave speed:

$$h \ll D/v: \ C_{\text{th}} = \frac{a\rho_0 D}{hv^2(1 + u/v)} \tag{S82}$$

and

$$h \gg D/v: \ C_{\text{th}} = \frac{2a\rho_0}{\pi v(1 + u/v)}. \tag{S83}$$

For neutrophils and the information wave front presented in the main text (reproduced in Fig. 9), $1 + u/v \approx 1 + (0.3\ \mu\text{m/s})/(2\ \mu\text{m/s}) = 1.15$ and the effect of chemotaxis on the asymptotic dynamics is small.

#### Effect on transient dynamics

To study the effect of chemotaxis on the transient dynamics of neutrophil swarming, we utilize a modified version of our Green's function method reported in Section 9 to propagate (S78a) and (S78b). One can do so using the same algorithm described previously, but with the following modifications with $r_c(t)$ the information wave front:

- For radii $r$ at time $t$ satisfying $r_i < r < r_c(t)$, find the time $t^*(r,t)$ at which the neutrophils at radius $r$ at time $t$ began chemotaxing inward. This time satisfies the relationship $r_c[t^*] - r = u(t - t^*)$.

- The density at $r$ is therefore given by the ratio of $r_c[t^*]$ to $r$ with an additional factor of one plus the ratio of the inward advection speed to the outward-propagating wave speed at $r_c[t^*]$ and $t^*$:

$$\rho(r,t) = \rho_0 \frac{r_c[t^*]/r}{1 + u\left(\frac{\partial r_c[t^*]}{\partial t}\right)^{-1}}. \tag{S84}$$

  This is analogous to (S79) and reduces to (S79) in the asymptotic limit of $r_c(t) = vt$.

- We assume that once the cells reach the target edge, they pack inward at a maximum density, in units of the cell diameter $d_c$, of $\rho_{\max} = 1/d_c^2$. For the experiments we discuss [8], this means that $\rho_{\max} \approx 10\ \rho_0$.

With all of these adjustments, and using the reported value of $u \approx 20\ \mu$m/s in [8], we arrive at the navy information wave front in Fig. 9A. This curve is a fit by eye to the experimental information wave front and has fit parameters of $D = 1.5 \times 10^{-10}$ m$^2$/s and $v = 1.73\ \mu$m/s, corresponding to a threshold concentration of $C_{\text{th}}/a\rho_0 = \frac{2}{\pi v(1+u/v)} \approx 3.07 \times 10^5$ s/m. For reference, the black curve in Fig. 9A is the information wave front from Fig. 4 of the main text, for which $D = 1.25 \times 10^{-10}$ m$^2$/s and $C_{\text{th}}/a\rho_0 = \frac{2}{\pi v(1+u/v)} \approx 3.67 \times 10^5$ s/m. Thus, including chemotaxis only negligibly affects our fit values.

To compare the two models in Fig. 9A, we plot both the concentration profile (Fig. 9B) and the concentration gradient (Fig. 9C) for a given critical radius (the dashed line in Fig. 9A). When one accounts for chemotaxis, the concentration profile near the target steepens relative to a model with a stationary cell distribution. We can therefore see that chemotaxis itself can lead to steeper concentration profiles, though the model we have explored here only accounts for an average inward drift of the cells.

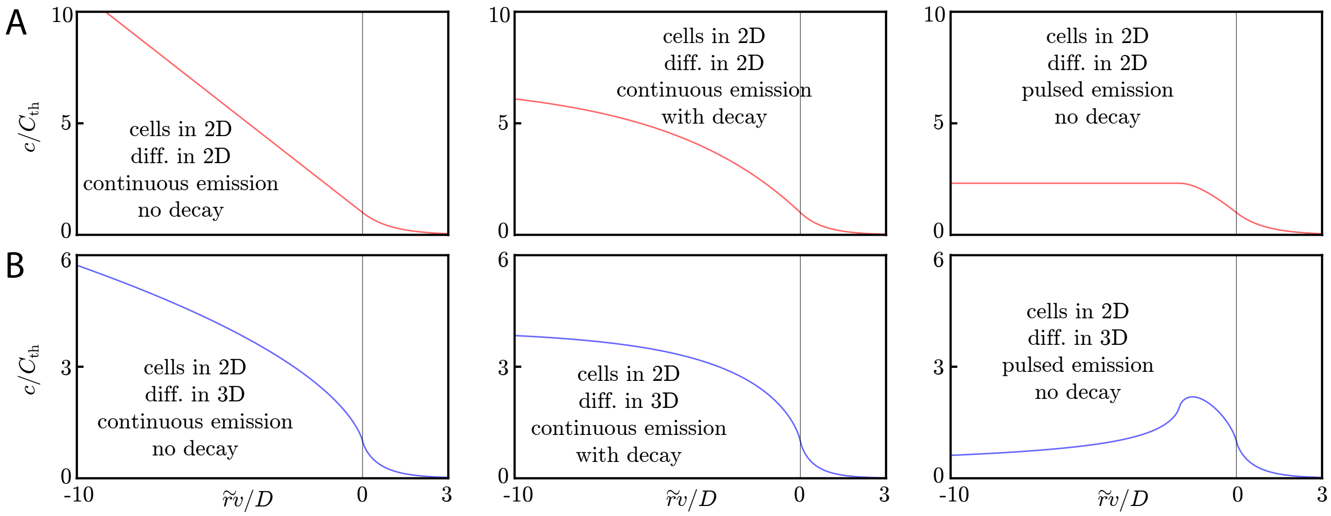


Figure 1: **A:** Signaling molecule concentration profiles for cells in 2D and diffusion in 2D. Here, we assume continuous emission and no signaling molecule decay (left panel), continuous emission and signaling molecule decay (middle panel), or pulsed emission with no signaling molecule decay (right panel). The decay constant for the middle panel is $\gamma = v^2/4D$ while the pulse width is $\tau = 2D/v$ in the right panel. For the left, center, and right panels, the threshold concentration is calculated according to (S11), (S27), and (S24), respectively. As discussed in the main text, the concentration profiles flatten (with respect to the profile generated by continuous emission without decay) inside the wave front once decay or pulsed emission is accounted for. **B:** Signaling molecule concentration profiles for the same cases as in **A**, but with diffusion in 3D. Compared to the case of continuous emission without decay, the concentration profiles flatten (when accounting for decay) or have a local maximum (in the case of pulsed emission).

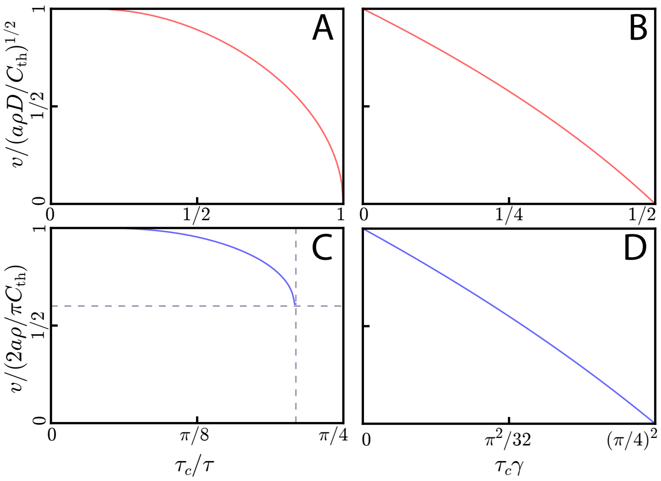


Figure 2: **A:** Wave speed $v$ for square pulse emission by cells in 2D with diffusion in 2D as a function of pulse width $\tau$. We normalize $v$ by the $\tau \to \infty$ wave speed of $(a\rho D/C_{\rm th})^{1/2}$. At $\tau = \tau_c = hC_{\rm th}/a\rho$, the wave speed goes to zero, and for shorter pulses there is no wave-like solution. **B:** Wave speed for continuous emission by cells in 2D with diffusion in 2D, accounting for signaling molecule decay at rate $\gamma$. For $\gamma = t_{\rm min,\ 2D} = 1/2\tau_c$, with $\tau_c$ as in panel **A**, the wave speed goes to zero and there are no wave-like solutions for larger decay rates. **C:** wave speed $v$ as a function of pulse width $\tau$ for square pulse emission of a signaling molecule by cells in 2D with a 3D diffusive environment. At $\tau \approx 1.53\tau_c = 1.53D(2C_{\rm th}/\pi a\rho)^2$ (vertical dashed line), there is a minimum wave speed of $v \approx 0.6 \times (2a\rho/\pi C_{\rm th})$ (horizontal dashed line), unlike with 2D diffusion. Importantly, $\tau \approx 1.53\tau_c$ is longer than the minimum initiation time of $4\tau_c/\pi$. Thus, for values of $\tau$ between these two, cells can reach the threshold concentration but cannot propagate a traveling information wave. **D:** Wave speed $v$ as a function of decay rate $\gamma$ for cells in 2D with diffusion in 3D. At $\gamma = (\pi/4)^2/\tau_c$, with $\tau_c$ as in **C**, the wave speed goes to zero.

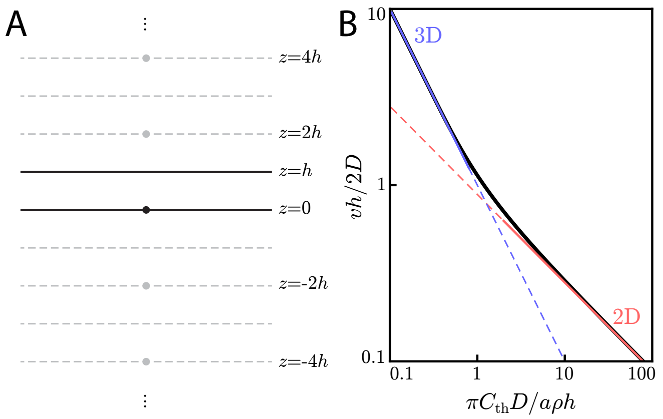


Figure 3: **A:** Method of images configuration for solving for concentrations in a finite-sized extracellular medium of thickness $h$. A point-like source (a cell) at $z = 0$ is placed in an extracellular medium of thickness $h$. In order to calculate the concentration with the appropriate boundary conditions of $\partial c/\partial z$ at $z = 0, h$, one need only add the contributions from "image cells" at $z = \pm 2jh$ for $j = 1, 2, \ldots$. **B:** Universal curve (black line) showing non-dimensionalized wave speed ($vh/2D$) versus non-dimensionalized threshold concentration ($\pi C_{\mathrm{th}} D/a\rho h$). In the limit of $vh/2D \gg 1$, we recover the familiar 3D scaling law of (S16) (blue line). In the limit of $vh/2D \ll 1$, we get the 2D scaling law of (S11) (red line).

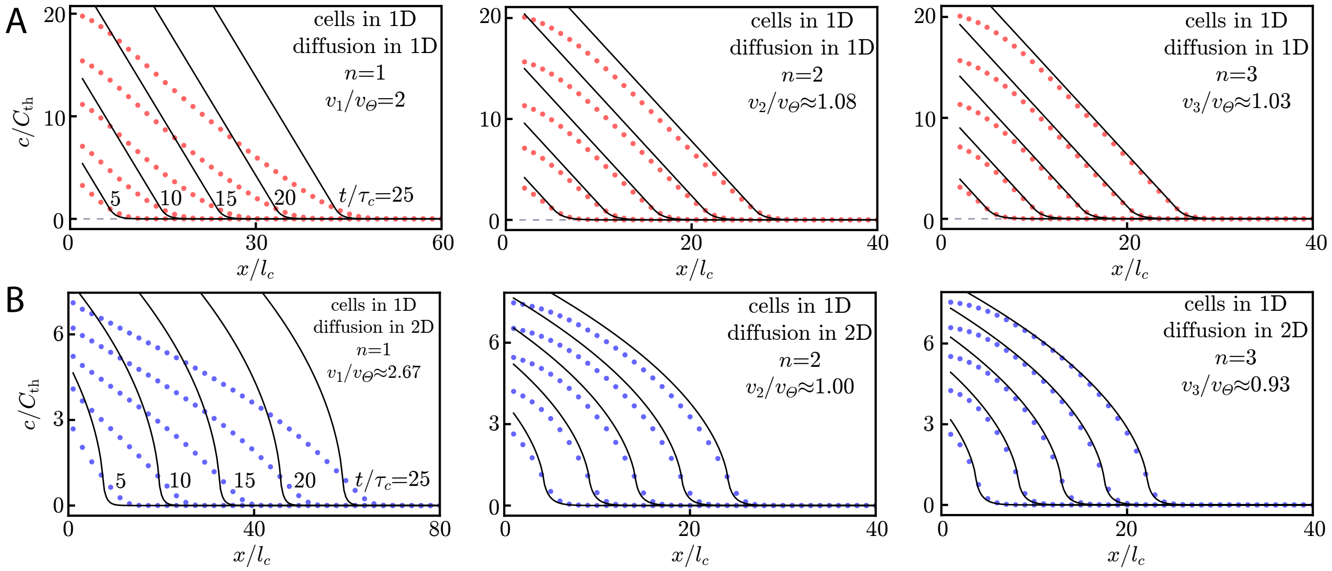


Figure 4: Wave speeds and profiles with Hill function activation. **A:** Numerical simulation of cells in 1D with diffusion in 1D and Hill function activation. The details of the simulation are described in the Supplementary Information text. For $n \geq 2$, we see good agreement between the Heaviside theory (black lines) and the Hill function numerics (red dots). Snapshots are shown at $t/\tau_c = 5, 10, 15, 20, 25$ where $\tau_c = h^2 C_{\mathrm{th}}/a\rho$ equals $D/v_\Theta^2$, the characteristic time scale for Heaviside activation. Note that the x-axis is scaled by the characteristic length $l_c = h(DC_{\mathrm{th}}/a\rho)^{1/2}$, which is the length scale for Heaviside activation with a delta function source, $D/v_\Theta$. In the insets, we display the wave speed for the order-$n$ Hill function, $v_n$, compared to the Heaviside activation wave speed, $v_\Theta = (a\rho D/h^2 C_{\mathrm{th}})^{1/2}$. Our fit to the $n = 1$ data gives $v_{n=1} \approx 1.93 v_\Theta$, but we have shown in Section 5 that $v_{n=1} = 2v_\Theta$, meaning that the wave speeds in the insets are slight underestimates. **B:** Numerical simulation of cells in 1D and diffusion in 2D with Hill function activation. For $n \geq 2$, the wave speed and concentration profiles (blue dots) agree well with the Heaviside theory (black lines). The theory plotted here assumes a delta-function-like source with respect to the extra diffusive dimension, as in (S12). The numerics, however, use a very narrow ($H = l_c/10$) Gaussian source. Here, the characteristic length $l_c = \pi h D C_{\mathrm{th}}/2a\rho$ equals $D/v_\Theta$, the length scale for Heaviside activation. The x-axis is scaled by the same quantity. Snapshots are shown at $t/\tau_c = 5, 10, 15, 20, 25$ with the characteristic time $\tau_c = D(\pi h C_{\mathrm{th}}/2a\rho)^2$ equal to $D/v_\Theta^2$, the time scale for Heaviside activation. In the insets, we display the wave speed for the order-$n$ Hill function, $v_n$, compared to the Heaviside activation wave speed, $v_\Theta = 2a\rho/\pi C_{\mathrm{th}}$.

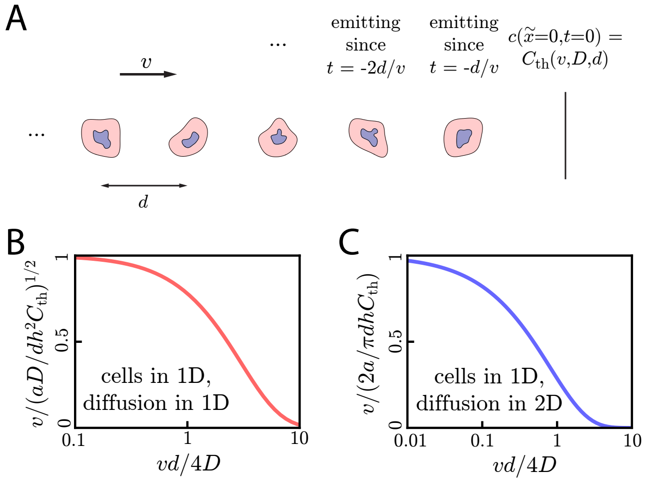


Figure 5: **A:** Cartoon showing the discrete lattice model of cell signaling relays for cells in 1D. Here, a concentration wave propagates rightward at speed $v$ through a group of cells spaced by a constant distance $d$. The concentration at $\tilde{x} = t = 0$ defines the threshold concentration in this discrete system and is a function of $v$, $D$, and $d$: $C_{\rm th} = C_{\rm th}(v, D, d)$. **B:** Wave speed $v$ compared to the continuous theory value of $\left(aD/dh^2C_{\rm th}\right)^{1/2}$ as a function of $vd/4D$ for cells in 1D with diffusion in 1D. We can see that the wave speed approaches the continuous theory value for $vd/4D \ll 1$. **C:** Same as **B** but for cells in 1D and diffusion in 2D.

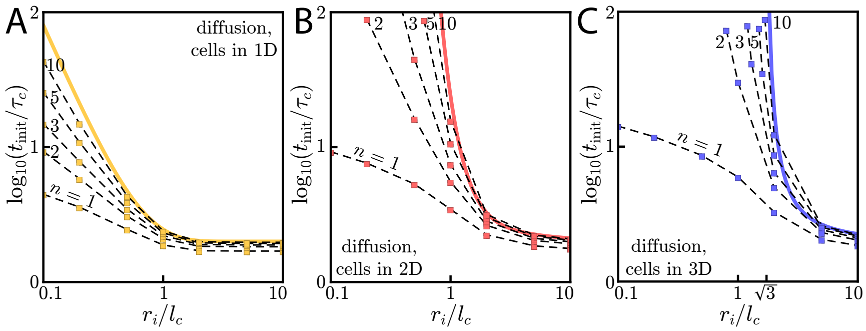


Figure 6: Diffusive wave initiation with Hill function activation. **A:** Wave initiation for cells in 1D with diffusion in 1D. Here, the initiation time $t_{\text{init}}$ is normalized by the characteristic time scale $\tau_c = h^2 C_{\text{th}}/a\rho$ and the initial signaling colony size $r_i$ is normalized by the characteristic length scale $l_c = \left(h^2 C_{\text{th}} D/a\rho\right)^{1/2}$. Yellow data points are individual simulations with dashed black guides to the eye connecting the yellow points. The solid yellow line corresponds to Heaviside-like activation and is reproduced from the main text. **B:** Same as **A**, but for cells in 2D and diffusion in 2D. Here, $\tau_c = hC_{\text{th}}/a\rho$ and $l_c = (hC_{\text{th}}D/a\rho)^{1/2}$. **C:** Same as **A**, but for cells in 3D and diffusion in 3D. Here, $\tau_c = C_{\text{th}}/a\rho$ and $l_c = (C_{\text{th}}D/a\rho)^{1/2}$. We note the value of $r_i = \sqrt{3}l_c$, the value below which waves fail to initiate for Heaviside-like activation.

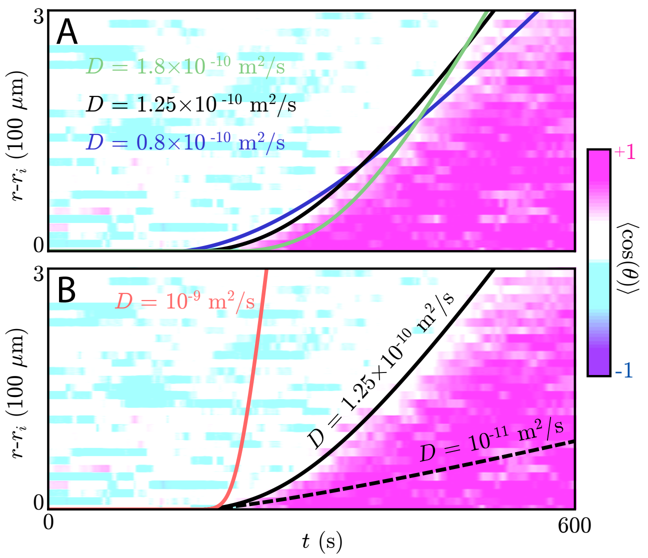


Figure 7: **A:** Plot showing the information waves for various choices of diffusion constant (black) overlaid on the experimental chemotactic index data from Reategui *et al.* (color plot) [8]. Here, we take $r_i = 100\ \mu$m, though the target in the experiment is a smaller, oblong object. The size of the target has no effect on the convex shape of the information wave front. The black line is reproduced from the main text and has $D = 1.25 \times 10^{-10}$ m$^2$/s and asymptotic wave speed $v \approx 1.7\ \mu$m/s (the threshold concentration is $C_{\rm th}/a\rho = 2/\pi v$). This diffusion constant is consistent with a small molecule like LTB4, and the resulting information wave dynamics can be made to fit the information wave front – both the initiation time of $\approx 200$ s and the transient dynamics. Other choices of parameters (green: $D = 1.8 \times 10^{-10}$ m$^2$/s, $v \approx 2.3\ \mu$m/s, navy: $D = 0.8 \times 10^{-10}$ m$^2$/s, $v \approx 1.3\ \mu$m/s) give information wave fronts that are also roughly consistent with the experimental data. **B:** However, with a much larger ($D = 10^{-9}$ m$^2$/s, red) or smaller ($D = 10^{-11}$ m$^2$/s, dashed) diffusion constant, an information wave with the correct initiation time does not have the correct transient dynamics. The wave speeds for these larger and smaller diffusion constants are, respectively, $v \approx 11\ \mu$m/s and $v \approx 0.3\ \mu$m/s.

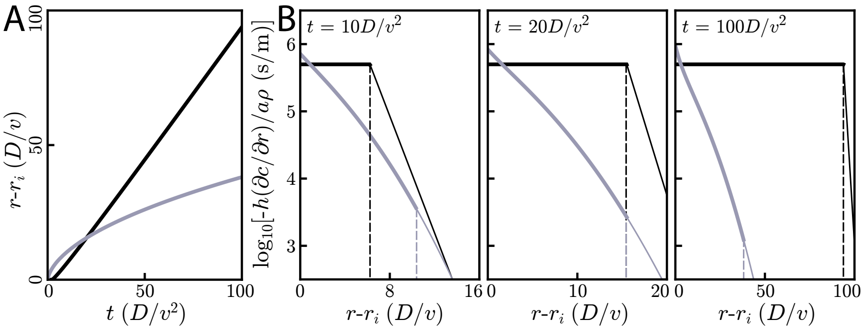


Figure 8: Comparison of relay and simple diffusion for cells and diffusion in 2D. All plots are for $D = 10^{-10}$ m$^2$/s, $v = 2$ $\mu$m/s, and $r_i = 4D/v$. **A:** Information fronts for relay (black) and simple diffusion (grey) models. The information wave travels like $\sqrt{t}$ for simple diffusion and $vt$ for the relay. The threshold concentration for the simple diffusion model is chosen such that the its information front intersects the relay model at $t = 20D/v^2$. Thus, $hC_{\mathrm{th,relay}}/a\rho = 25$ s while $hC_{\mathrm{th,diff.}}/a\rho \approx 0.25$ s. **B:** Snapshots of radial gradients at various times for both the relay (black) and simple diffusion (grey) models. The dashed vertical lines indicate the location of the wave fronts. For short times (left) the relay's information wave front lags behind the simple diffusion model's information front, though it later catches up (middle) and passes it (right). At all times, for cells just inside the wave front, the relay model creates gradients that are orders of magnitude larger than does simple diffusion.

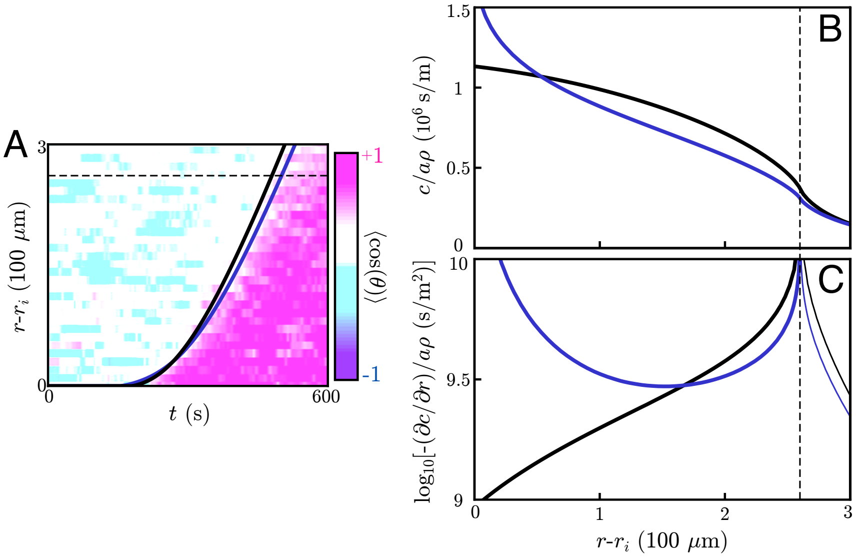


Figure 9: **A:** Information wave fronts for cell signaling relays with (navy) and without (black) chemotaxis. The information wave fronts are overlaid on the experimental chemotactic index data from Reategui *et al.* (color plot) [8]. The black curve is reproduced from the main text. Both models can account for the observed information wave fronts by fitting two parameters: the signaling molecule diffusivity, $D$ and the threshold concentration, $C_{\mathrm{th}}$. **B/C:** Concentration profiles (**B**) and gradients (**C**) generated by the signaling relay models in **A**. The wave front is indicated in all panels by the dashed line, and the concentration profiles and gradients are plotted at the times such that the threshold concentration is equal to the concentration at the wave front. When one accounts for chemotaxis, the concentration profiles near the target steepen relative to models without chemotaxis.
